# Modelling APOL1-mediated kidney inflammation and fibrosis using a partially reprogrammed urine derived SIX2-positive renal progenitor cell line

**DOI:** 10.1101/2025.07.28.667124

**Authors:** Chantelle Thimm, Rosanne Mack, Osmond Adjei-Aruna, Wasco Wruck, James Adjaye

## Abstract

**Background:** CKD affects approximately 850 million people worldwide and is a leading cause of mortality. Podocytes, cells in the kidney are terminally differentiated and incapable of division *in vivo* making the establishment of primary cultures particularly challenging. The ability of cells to proliferate and avoid senescence is closely linked to telomere length. When telomere length becomes critically reduced, it results in cellular senescence.

**Methods:** We present the successful rejuvenation of a human SIX2-positive renal progenitor cell line derived from the urine of a 30-year-old West African male (UM30-OSN). To achieve partial reprogramming, plasmids expressing the Yamanaka factors OCT4, SOX2, NANOG, c-Myc, and KLF4 were employed.

**Results:** UM30-OSN expresses the pluripotency-associated marker SSEA4, renal stem cell markers such as SIX2, CD133 and CD24, determined by immunofluorescence, FACS and qPCR. Expression analysis revealed downregulation of senescence markers p21and p53 and upregulation of proliferation-associated genes PCNA, KI67 and TERT, confirming rejuvenation.

Upon podocyte differentiation, UM30-OSN cells expressed podocyte-specific markers NPHS1, NPHS2, SYNPO and CD2AP. Comparative transcriptome analyses revealed a correlation co-efficiency (R^2^ = 0.88) with the immortal podocyte line AB 8/13.

To demonstrate the usefulness of UM30-OSN to model APOL1-mediated kidney disease, we investigated the effects of Interferon-γ (IFN-γ) on UM30-OSN derived podocytes and evaluated the potential of the JAK1/JAK2 inhibitor Baricitinib to mitigate IFN-γ–induced cellular responses. IFN-γ stimulation resulting in increased phosphorylation of STAT1, activation of APOL1, upregulation of pro-inflammatory and fibrotic markers such as, IL-6, TGF-β, Vimentin, Fibronectin, and morphological changes indicative of cell stress. Pre-treatment with Baricitinib effectively inhibited STAT1 phosphorylation, reduced expression of pro-inflammatory and fibrosis-associated genes, and preserved podocyte morphology.

**Conclusion:** Given their robust proliferation capacity, UM30-OSN cells represent a valuable additional model for investigating kidney-associated diseases such the contribution of APOL1 high-risk variants to kidney injury and fibrosis.

## Background

Kidneys comprise approximately one million nephrons, which play vital roles in regulating systemic pH, maintaining fluid homeostasis, filtration of blood and electrolyte balance. Each nephron consists of a renal corpuscle, which includes the glomerular filtration barrier and a complex renal tubular system. The glomerular filtration barrier is formed by podocytes, which are highly specialized epithelial cells that envelop the glomerular capillaries and extend into the urinary space of Bowman’s capsule. These cells possess a large cell body with numerous foot processes that interdigitate with those of adjacent podocytes. These lamellipodia-like extensions form intercellular junctions across the slit diaphragm, ensuring selective permeability based on molecular size during glomerular filtration. Several podocyte-specific genes have been identified, whose protein products are critical for the integrity and function of the filtration barrier. These include NPHS2 (Podocin), NPHS1 (Nephrin), CD2AP (CD2-associated protein) and SYNPO (Synaptopodin).

Currently, animal models, including rats and mice, are extensively utilized in the study of kidney diseases.

However, these models exhibit significant anatomical, histological, and molecular differences compared to human kidneys. This discrepancy underscores the urgent need for more physiologically relevant human experimental models.

To date, the use of induced pluripotent stem cells (iPSCs), and the temperature-sensitive SV40 conditionally immortalized cell lines has become widespread in kidney-associated research ^1–4^.

The growth-promoting effect of the SV40 DNA sequence was first reported in 1965 ^5^. This immortalized ability was first described in detail by Chang et al. in 1984 ^6^, for which the tumor (T) antigen and the small T antigen appear to be responsible. The small T antigen facilitates the replication of the viral genome and can accelerate the G1 and S phases of the cell cycle. In parallel, the T antigen interacts with the tumor suppressor retinoblastoma family (pRB) and p53 ^7^.

In earlier studies have shown that it is possible to rejuvenate adult somatic cells by transient over expression of the Yamanaka factors without the sole dependence on SV40 T antigen ^8^. Interferon-gamma (IFN-γ) acts as a pivotal pro-inflammatory cytokine implicated in immune-mediated kidney diseases. It activates JAK/STAT signalling, inducing transcription of inflammatory and pro-fibrotic genes such as APOL1, which is strongly associated with increase the risk of kidney damage, including fibrosis, leading to chronic kidney disease (CKD) and potentially kidney failure in individuals of West African descent carrying the G1 or G2 risk variants ^9–11^. Functional investigation of APOL1 has largely relied on over-expression models in mice and HEK293 cells to over-express APOL1-G0, -G1, or -G2 variants. These studies revealed mitochondrial dysfunction, altered gene expression profiles, and impaired respiration in HEK cells over-expressing G1 or G2 variants ^11–13^. Although these models provide valuable mechanistic insights, they lack the characteristic features of human podocytes and the native regulatory context of APOL1.

APOL1 gene variants are associated with an increased risk of kidney disease, particularly in individuals of African origin. These variants can lead to various forms of kidney disease, including focal segmental glomerulosclerosis (FSGS), collapsing glomerulopathy, hypertensive nephropathy, CKD and end-stage kidney failure ^14–18^. The development of renal fibrosis characterizes CKD. Regardless of the primary cause, renal fibrosis is the hallmark of progressive CKD^19^. Fibrogenesis is initiated in response to chronic kidney damage; following renal injury, profibrotic factors are produced by injured tubular epithelial cells and infiltrating inflammatory cells that stimulate signaling events leading to ECM production. APOL1 risk variants are linked to accelerated fibrosis in the kidneys, contributing to the progression of kidney disease^20^, which has been shown in human cellular^21^ and mouse models^22^. In a transgenic mouse model expressing the podocyte-specific APOL1-G2 risk variant (G2APOL1), these variants led to activation of STING and NLRP3 inflammasomes. Triggered pronounced renal fibrosis, increased albuminuria and impaired renal function^22^.

In the present study, we used rejuvenated SIX2-positive renal progenitor cells derived from the urine of a male donor of West African ancestry, differentiated into podocytes (UM30-OSN). This model combines genetic authenticity with the physiological relevance of podocytes and enables, for the first time, the investigation of IFN-γ–induced APOL1 regulation and the mechanisms of Baricitinib action in a more disease-relevant cellular context.

These cells exhibit stable morphology and proliferation capacity *in vitro* for over one year of culture. The UM30-OSN cell line expresses the homeodomain transcriptional regulator SIX2, representing a multipotent, self-renewing progenitor population capable of differentiating into all nephron cell types ^23,24^. We further analyzed gene and protein expression, morphological cellular changes, and activation of the JAK/STAT signaling pathway, as well as the ability of Baricitinib to suppress these IFN-γ–mediated effects.

Due to their high proliferation capacity and ease of handling, our immortalized male UM30-OSN cell line of West African origin, which we believe is one of its kind, provides a unique tool for studying the molecular basis of APOL1-mediated inflammation and fibrosis in the kidney.

## Methods

### Cell Culture Conditions

Urine-derived renal progenitor cells (UdRPCs) of a 30-year old man (UM30) were isolated and maintained in proliferation medium (PM) as described in Rahman et al ^25^. Furthermore, approximately 250,000 SIX2-positive urine-derived cells from UM30 were immortalized by nucleofection of an episomal not integrative plasmid-based ectopic expression of Yamanaka reprogramming factors co-expressed SV40 large T antigen, which inactivates the tumor suppressors pRB and p53 (UM30-OSN)^26,27^. For more efficiency, pathway inhibitions: TGFβ /SB431245 (Stemgent #04-0014), MEK/PD0325901 (Stemgent #04-0006) and GSK3β/CHIR99021 (Stemgent #04-0004) were added. Cells were cultured for 24 h in PM supplemented with 10 μM Y27632 on Matrigel® at 37 ◦C and 5 % CO2. After 24 h, the medium was changed to StemMACS iPS-Brew XF Medium (Miltenyi Biotec, Bergisch Gladbach,Germany) with daily medium changes. Besides the well-defined iPSC colonies clusters of partially reprogrammed cells were identified. These rapidly proliferating cells are still mesenchymal unlike iPSCs and they resemble the primary urine cells. These clusters were picked and expanded on plastic, cultured at 37 °C, 5% CO2. The cells were cultured in proliferation medium (PM) composed of 50% DMEM high Glucose (Gibco®, life technologies, California) and 50% Keratinocyte growth basal medium (Lonza, Basel, Switzerland) supplemented with 5 ng/ml bFGF. The DMEM high Glucose medium is supplemented with 10% fetal bovine serum (Gibco®), 0.1% Non-Essential Amino Acid (Gibco®), 0.5% Glutamax (Gibco®) and 1% Penicillin and Streptomycin (Gibco®). Differentiation into podocytes followed our previously published protocols ^2,28,29^. In brief, 25,000 cells per 6-well and 100,000 cells per 12-well were first cultured for 24 h in PM on Corning® Collagen I (Merck, Darmstadt, Deutschland) coated plates till approximately 70% confluent. On the next day, the medium was replaced with Advanced RPMI 1640 (Gibco®) supplemented with 10% fetal bovine serum, 1% Penicillin and Streptomycin, and 30 μM retinoic acid (Sigma-Aldrich Chemistry, Steinheim, Germany). After seven days the typical podocyte morphology was observed.

We used the already established human immortal cell line (AB 8/13) described by Saleem et. al. as a reference cell line ^3^. The human immortal cell line are expanded at 33°C, 5% CO2 on plastic. The cells were cultured in RPMI 1640 (Gibco®, life technologies, California) supplemented with 10% fetal bovine serum (Gibco®,) and 1% Penicillin and Streptomycin (Gibco®,). For further differentiation into podocytes, the cells were seeded on Corning® Collagen I (Merck, Darmstadt, Deutschland) coated plates. Both UM30-OSN and AB 8/13 were cultured in podocyte differentiation medium-Advanced RPMI 1640 (Gibco®) supplemented with 10% fetal bovine serum, 1% Penicillin and Streptomycin, and 30 μM retinoic acid (Sigma-Aldrich Chemistry, Steinheim, Germany) at 37 °C, 5% CO2 for approximately 14 days. The medium was changed every third day.

To investigate the effect of IFN-γ on UM30-OSN podocytes, we pre-treated them for 24 h with or without 1 µM Baricitinib (Medchemexpress #HY-15315) followed by an additional 24 h treatment of 100 ng/ml IFN-γ (Thermofisher #300-02-100UG).

### RNA isolation

For RNA isolation from the various cell lines, the ZYMO Research Kit Direct-zol™ RNA Miniprep R20 was utilized following the manufacturer’s protocol. Cells were detached by adding Tryple E and incubated for 5 min. Afterwards, a centrifuging step was added 4 mins 1000 g. Cell pellet was resuspend in 350 μl volume of 95–100% ethanol (EtOH) and thoroughly mixed by pipetting. The resulting solution was transferred to a Zymo-Spin™ column and centrifuged at 12,000 g for 30 seconds at 4 °C. To degrade genomic DNA, the column was washed with RNA Wash Buffer and centrifuged again at 12,000 g for 30 seconds at 4 °C. Subsequently, the column was treated with DNA digestion mix (1:16 dilution of DNase I in DNA Digestion Buffer) for 15 minutes. Following digestion, the column was rinsed twice with RNA Prewash Buffer, each followed by centrifugation at 12,000 g for 30 seconds at 4 °C. A final wash with RNA Wash Buffer was performed, including a 2-minute centrifugation step. After drying the column by centrifuging for 1 minute, it was transferred to a fresh RNase-free Eppendorf tube. RNA was eluted by adding 35 μL of DNase/RNase-free water to the column and centrifuging for 1 minute at 16,000 g. The eluted RNA samples were immediately placed on ice, and concentrations measured with a Nanodrop device.

### Relative Quantification of podocyte-associated gene expression by real-time PCR

This followed our previously published protocols^2,28,29^. In brief, Quantitative real-time PCR of podocyte-associated gene expression was performed as follows: Samples were run in triplicate on a 384-well reaction plate and Power Sybr Green PCR Master Mix (Applied Biosystems, Foster City, CA, USA) using Step One Plus Real-Time PCR systems. The amplification conditions were denaturation at 95 ^◦^C for 13 min followed by 37 cycles of 95 ° C for 50 s, 60 ^◦^C for 45 s, and 72 ^◦^C for 30 s. Primer sequences are listed in supplemental table S2. Results were analyzed by the 2^−ΔΔ-CT^ method and specified in the fold change expression.

### FACS

UM30-OSN cells were washed with PBS and incubated for approximately 5 min at 37 °C, 5% CO2 in 0.25% Trypsin EDTA. The reaction was stopped with 1:2 DMEM and the cell suspension was transferred to a 15 ml Falcon. This was followed by centrifugation at 1000x g for 4 min. The resulting cell pellet was resuspended with 1 – 2 ml 3% BSA and was blocked for 20 min. at room temperature. Cells were then separated using a 0.4 μM mesh strainer (Thermo Fisher Scientific, Massachusetts, USA) and placed in FACS tubes (Corning, New York, USA,). Afterward 3 ml of MACS buffer (0.5 % BSA, 2 mM EDTA, PBS) were added to each tube followed by an additional centrifugation step. The cells were resuspended at up to 1×10^7^ cells / ml in 4% PFA (Polysciences, Warrington, FL, USA) and incubated for 20 min at room temperature. Followed by two washing steps with PBS. After each washing step the cells were centrifuged at 1000x g for 10 min. Afterwards the cells were diluted in 98 μl MACS Buffer and 2 μl of the primary antibody CD133 (Thermo Fisher Scientific, Massachusetts, USA), CD24 (Thermo Fisher Scientific, Massachusetts, USA) and SIX2 (Abnova, Heidelberg, Germany). The antibody was incubated overnight at 4 °C. Cells were washed by adding 1 – 2 ml MACS Buffer and centrifuged at 1000x g for 10 min. The supernatant was completely aspirated followed by 1 h incubation of the secondary antibodies at room temperature in the dark. 0.2 μl of the secondary antibody Alexa 555 and 647 (Thermo Fisher Scientific, Massachusetts, USA) was diluted in 100 μl MACS Buffer for each tube. The cells were washed once with 0.05% Tween-20 diluted in PBS. The cell pellets were then resuspended in a suitable amount of MACS Buffer Measurement of fluorescence intensity was demonstrated using Beckman Gallios Flow Cytometer (Beckman Coulter). In each case, 30,000 cells were measured, and data were analyzed using FlowJo software (BD Biosciences). Detailed information on the antibodies used can be found in supplementary table 1.

### Immunofluorescence-based staining

For the immunocytochemistry, urine-derived cells were fixed with 4% paraformaldehyde (PFA) (Polysciences, Warrington, FL, USA). Followed by three wash steps with PBS for 5 min each. To block unspecific binding sites, the fixed cells were incubated with a blocking buffer containing 10% normal goat or donkey serum, 1% BSA, and 0.5% Triton X-100. (Sigma-Aldrich Chemistry,) and 0.05% Tween-20 (Sigma-Aldrich Chemistry,) for 2 h at room temperature. The primary antibodies were diluted in blocking buffer according to supplemental table 1 and incubated at 4 °C overnight. The following day, the cells were washed once with 0.05% Tween-20 diluted in PBS and two times with PBS only. After washing them, the fluorochrome-conjugated secondary antibody and HOECHST (1:5000) (Thermo Fisher Scientific, Waltham, MA, USA), were added and incubated for 1 h at room temperature protected from light. The secondary antibodies were diluted in blocking buffer according to supplemental table 1. Visualization of cells employed a fluorescence microscope (LSM700).

### Western Blot Analysis

Podocytes were lysed in RIPA buffer (Sigma-Aldrich Chemistry) containing 5 M NaCl, 1% NP-40, 0.5% DOC, 0.1% SDS, 1 mM EDTA, 50 mM Tris, pH 8.0, and freshly added 10 μL/mL protease- and phosphatase inhibitor (Sigma-Aldrich). 20 μg of each obtained protein lysate was dissolved in a 10% sodium dodecyl sulfate PAGE gel and transferred to an Immobilon-P membrane (Merck Millipore, Burlington, VT, USA). The membranes were incubated with the primary antibody overnight at 4 °C. Then washed three times with 0.1% Tween-20 in Tris-buffered saline followed by 1 h incubation of the secondary antibodies at room temperature. For visualization of the blotted proteins the Pierce^TM^ ECL Western Blotting Substrate solutions from Thermo Fisher were used (Thermo Fisher Scientific, Massachusetts, USA). were used. Both solutions were diluted 1:1. Quantification was carried out with ImageJ. Detailed information on the antibodies used can be found in the supplementary table 1.

### Microarrayrray and bulk NGS analysis

For the microarray and NGS experiments, 1 μg RNA preparations were hybridized on the Human Clariom S Gene Expression Array (Affymetrix, Thermo Fisher Scientific, Waltham, MA, USA) and were used for a 3’RNA-Seq assay on an Illumina NextSeq2000 sequencing system at the core facility Biomedizinisches Forschungszentrum (BMFZ) of Heinrich Heine University-Düsseldorf.

The fastq files containing the RNA-seq raw data were aligned against the GRCh38 genome via the software HISAT2 ^30^ using the parametrization suggested by Barruzzo et al. ^31^: hisat2 -p 7 -N 1 -L 20 -i S,1,0.5 -D 25 -R 5 --mp 1,0 --sp 3,0 -x hisatindex/grch38_r109 -U input.fastq.gz -S output.sam. The resulting BAM files were sorted using the SAMtools software ^32^ and the reads were condensed to counts per gene with the subread (1.6.1) featurecounts routine ^33^ using the ENSEMBL annotation file Homo_sapiens.GRCh38.109.gtf. The resulting table of counts per gene were imported into the R/Bioconductor ^34^ and normalized with the voom ^35^ algorithm from the limma package ^36^ filtering expressed genes (counts per million > 1 in at least one sample). Affymetrix raw data was normalized via the Robust Multi-array Average (RMA) method from the R/Bioconductor package oligo ^37^. Gene expression heatmaps were generated using the heatmap.2 function from the gplots package ^38^ employing Pearson correlation or Euclidean distance as distance measures. Gene expression was dissected in Venn diagrams drawn with the Venn Diagram package ^39^. Genes were considered expressed when their detection-p-value, calculated as described before ^40^, was less than 0.05 in microarrays or when there was more than 1 read in NGS data. The subsets of genes obtained from the Venn diagram analysis were analyzed for over-representation of gene ontologies (GOs) using the GOstats R package ^41^ and KEGG (Kyoto Encyclopedia of Genes and Genomes) pathways ^42^. The n=60 most significant pathways and GO terms (p < 0.05) were visualized via the R package ggplot2 ^43^ in dotplots indicating p-value, number and ratio of involved genes. Gene expression data will be available online in the National Centre of Biotechnology Information (NCBI) Gene Expression Omnibus.

### Ethical Statement

In this study, urine samples were collected with the informed consent of the donors and the written approval (Ethical approval Number: 2017-2457_3) of the ethical review board of the medical faculty of Heinrich Heine University, Düsseldorf, Germany. All methods were carried out in accordance with the approved guidelines. The medical faculty of Heinrich Heine University approved all experimental protocols. All patients provided written informed consent.

### Statistics

All samples were analyzed in triplicate. The mean value of these was used for the calculations. The statistical significance was calculated using the two-sample Student’s *t*-test with a significance threshold *p* = 0.05. Data regarding the Baricitinib and IFN-γ treatment were analyzed using one-way ANOVA to assess statistical significance between groups. A p-value of < 0.05 was considered statistically significant. All experiments were analyzed n=3.

## Results

### Successful immortalization of UM30-OSN

UM30-OSN cells were isolated directly from the urine of a 30-year-old West African male as described in Rahman et. al. ^25^. The cells were transfected with the episomal-based plasmids pEP4EO2SCK2MEN2L and pEP4EO2SET2K as in our previous iPSC lab resource publications ^27,44^ (Supplementary Figure S2).The vectors express SV40 and the reprogramming factors-OCT4, NANOG, c-MYC, KLF4 and SOX2. The immortalized cells are referred to as UM30-OSN, characterized by rapid proliferation than the primary UdRPCs, no morphological changes and full retention of the SIX2-positive mesenchymal progenitor cell characteristics. The proliferating cells were maintained for over a year without apparent morphological changes (Figure 1A). Because immortalization is associated with chromosomal alterations and genomic instabilities ^45^, we comparatively analysed the karyotype of UM30-OSN and the immortal podocyte cell line-AB 8/13 at the Institute of Human Genetics of the University Hospital Düsseldorf (Supplementary Figure S3).

**Figure 1:**
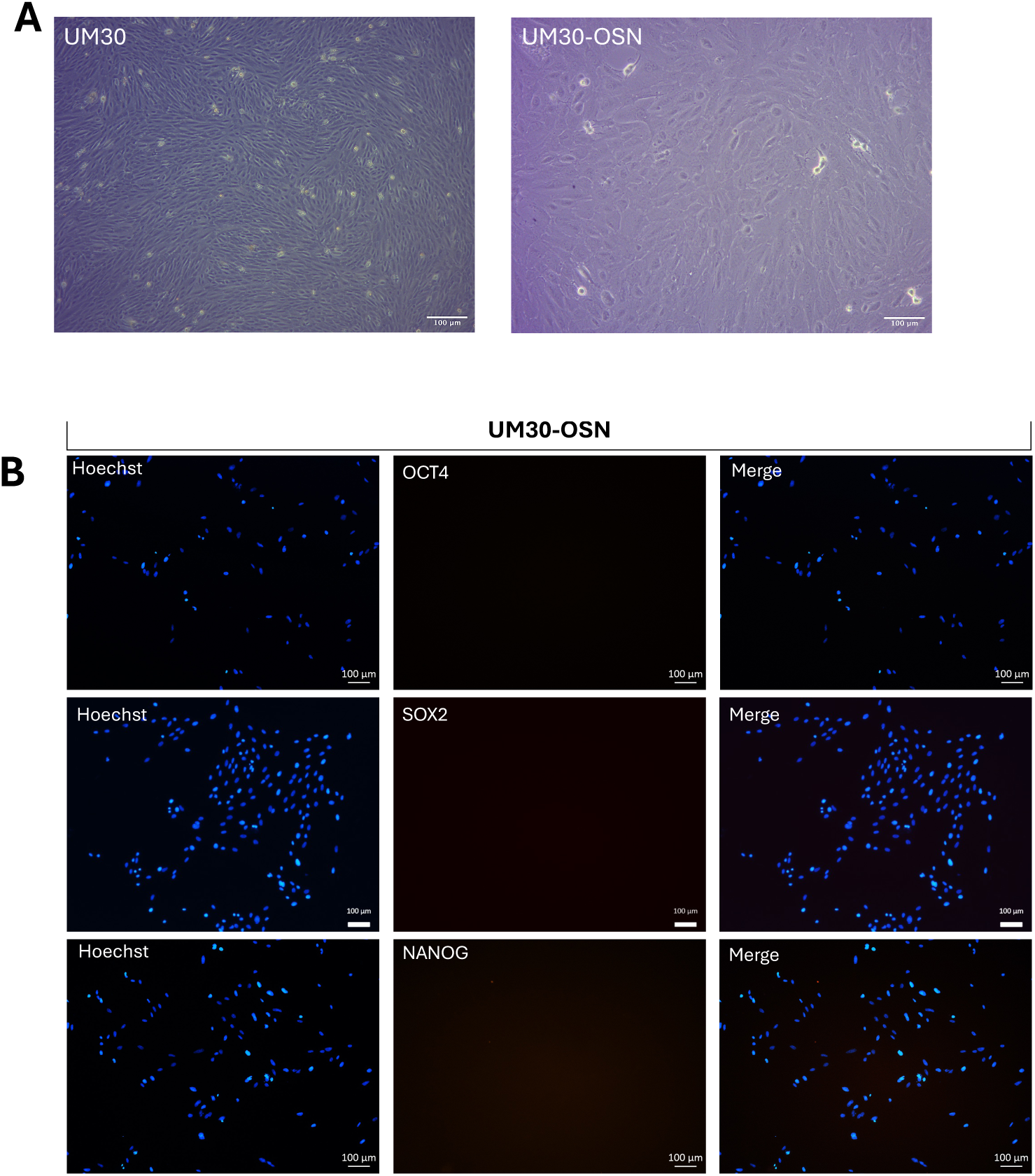
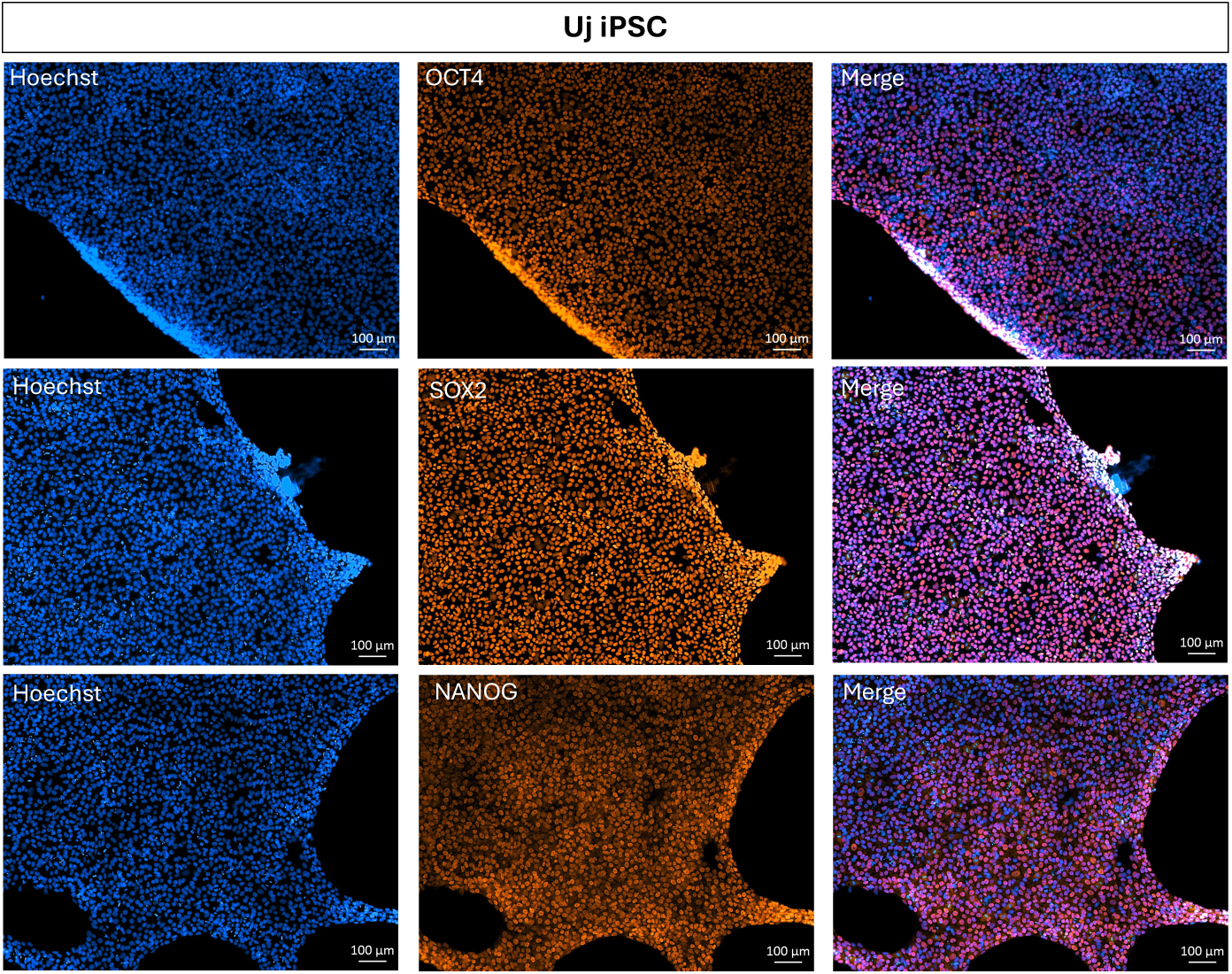
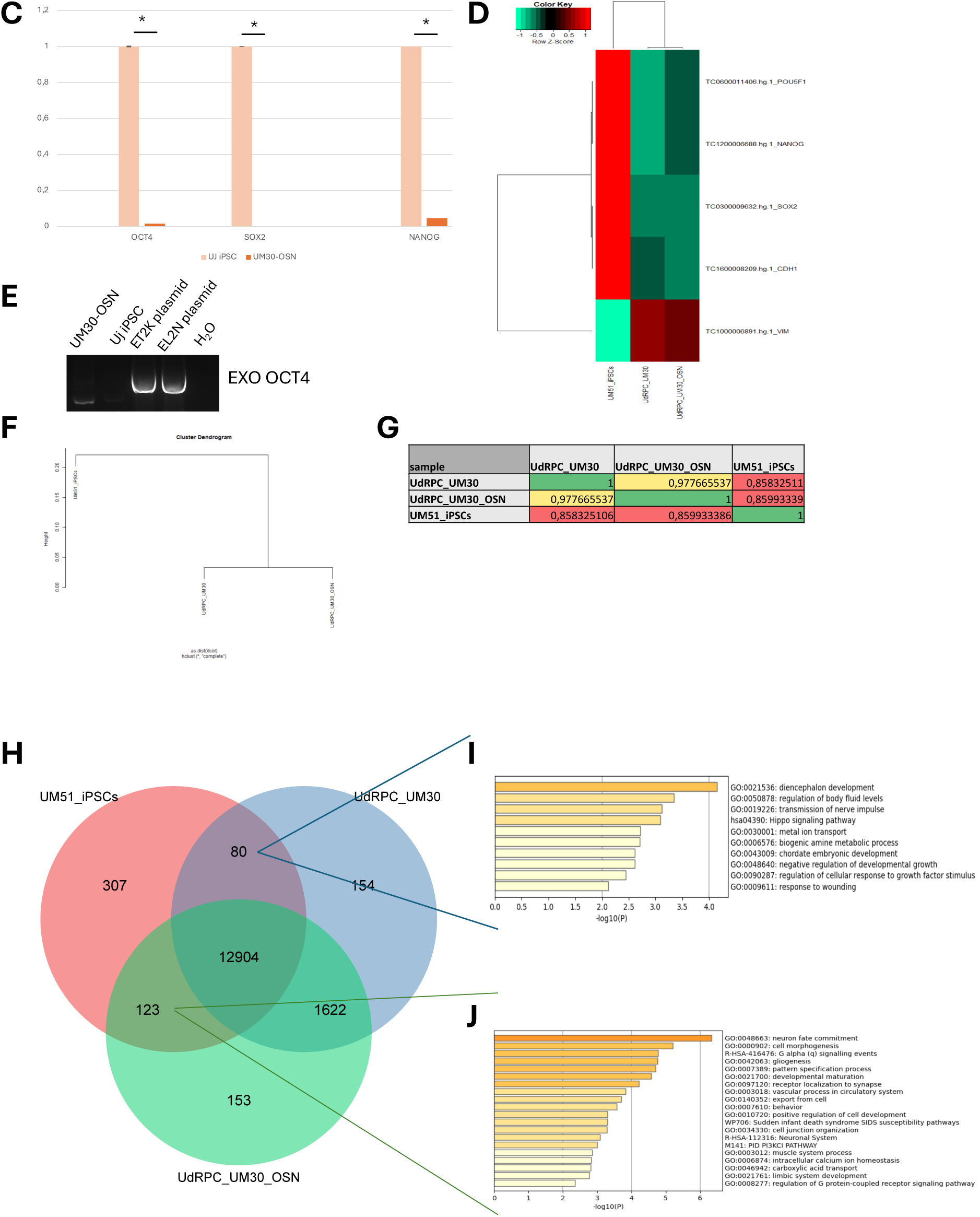
Successful immortalized UM30-OSN cell line derived from human urine. **(A)** The UM30-OSN cell line was partially reprogrammed using episomal plasmids encoding the Yamanaka factors. Light microscopy revealed morphological changes typical of early reprogramming stages. **(B)** The absence of full pluripotency was confirmed by immunofluorescence staining for OCT4, SOX2, and NANOG (scale bars: 100 μm). **(C)** qPCR analysis further validated the lack of expression of these pluripotency markers. **(D)** A heatmap displayed the expression levels of OCT4, SOX2, NANOG, E-CAD, and VIM, comparing UM30-OSN to a laboratory-derived induced pluripotent stem cell line (UM51-iPSC) used as a reference. **(E)** The absence of episomal plasmids was confirmed by semi-quantitative PCR targeting exogenous sequences. **(F–G)** A cluster dendrogram and correlation matrix demonstrated the transcriptional relationship between UM51-iPSC, UdRPC-UM30, and UM30-OSN**. (H)** Venn diagram-based transcriptome comparison was carried out between UM30-OSN, UdRPC UM30 and UM51 iPSC. **(I)** Metascape analysis of the 80 genes common expressed in the UM51 iPSC and UdRPC-UM30**. (J)** Metascape analysis carried out with the 123 genes exclusively expressed in the UM51-iPSC and UM30-OSN.

The primary cells-UM30 and the immortalized counterpart-UM30-OSN cell line grow as monolayers with typical rice grain morphologies (Figure 1A).

The absence of plasmids used in partial reprogramming was confirmed by immunofluorescence staining, quantitative real-time PCR, heatmap displayed the expression levels of OCT4, SOX2, NANOG and semi-quantitative PCR (Figure 1B). For comparison, an in-house induced pluripotent stem cell line (iPSC) UM51 iPSC was used as a control cell line to demonstrate that UM30-OSN is not pluripotent (Figure 1B), quantitative real-time PCR results (Figure 1C) and a heatmap of *OCT4, SOX2, NANOG, CDH1* and *VIM* (Figure 1D). For the semi-quantitative PCR, primers were designed, which amplified the exogenous OCT4 (Figure 1E).

Similar to the stable established cell line UM51 iPSC, the UM30-OSN cell line did not show a comparable band to the positive plasmid controls at 657 bp.

The immunocytochemistry-based expression detection was negative for OCT4, SOX2 and NANOG compared to the UM51 iPSC cell line (Figure 1B), therefore confirming the absence of the plasmids and therefore partial and not fully reprogrammed (Figure 1C). After the successful immortalization of UM30-OSN, we performed a comparative transcriptome analysis. Hierarchical clustering analysis comparing the transcriptomes of urine-derived renal progenitor UM30, the immortalized UM30-OSN cell line, and UM51 iPSC (Figure 1F). The UM50 iPSC cell line ^27^ serves as a direct comparison between a fully reprogrammed iPSC and the partially reprogrammed UM30-OSN. The correlation table indicates a positive relationship between UM30 and UM30-OSN (R^2^ = 0.90), which confirms the origin of the UM30-OSN (Figure 1G). The correlation co-efficient between UM30, UM30-OSN and UM51 iPSC is 0.85 (Figure 1G).

Additionally, a Venn diagram-based transcriptome comparison was carried out between UM30-OSN, UdRPC UM30 and UM51 iPSC (Figure 1H). A total 12904 genes were expressed in common expressed in all cell lines. Comparing the expressed genes (det-p < 0.05), 154 genes are exclusively expressed in UdRPC UM30, 153 genes in UM30-OSN and 307 genes in the UM51 iPSC (Figure 1H). Figure 1 represents a metascape analysis of the 80 genes only expressed in the UM51 iPSC and the UdRPC UM30, with GO:BP terms and pathways related to e.g. diencephalon development, regulation of body fluid levels and metal ion transport. The exclusive GO:BP terms and pathways related to the 123 genes exclusively expressed in the UM51 iPSC and UM30-OSN are e.g. cell morphogenesis, positive regulation of cell development and cell junction organization.

The full gene list of GOs and associated genes are in Supplementary table S4A.

### Characterization UM30-OSN

To determine the renal origin of UM30-OSN and thus its differentiation capacity, we performed immunofluorescence-based detection of the proteins, SSEA4 as a pluripotency marker, Vimentin as a cytoskeletal protein of mesenchymal cells, CD133 (Prominin-1) as a stem cell self-renewal marker and SIX2 as a multipotent self-renewing renal progenitor marker (Figure 2A).

**Figure 2:**
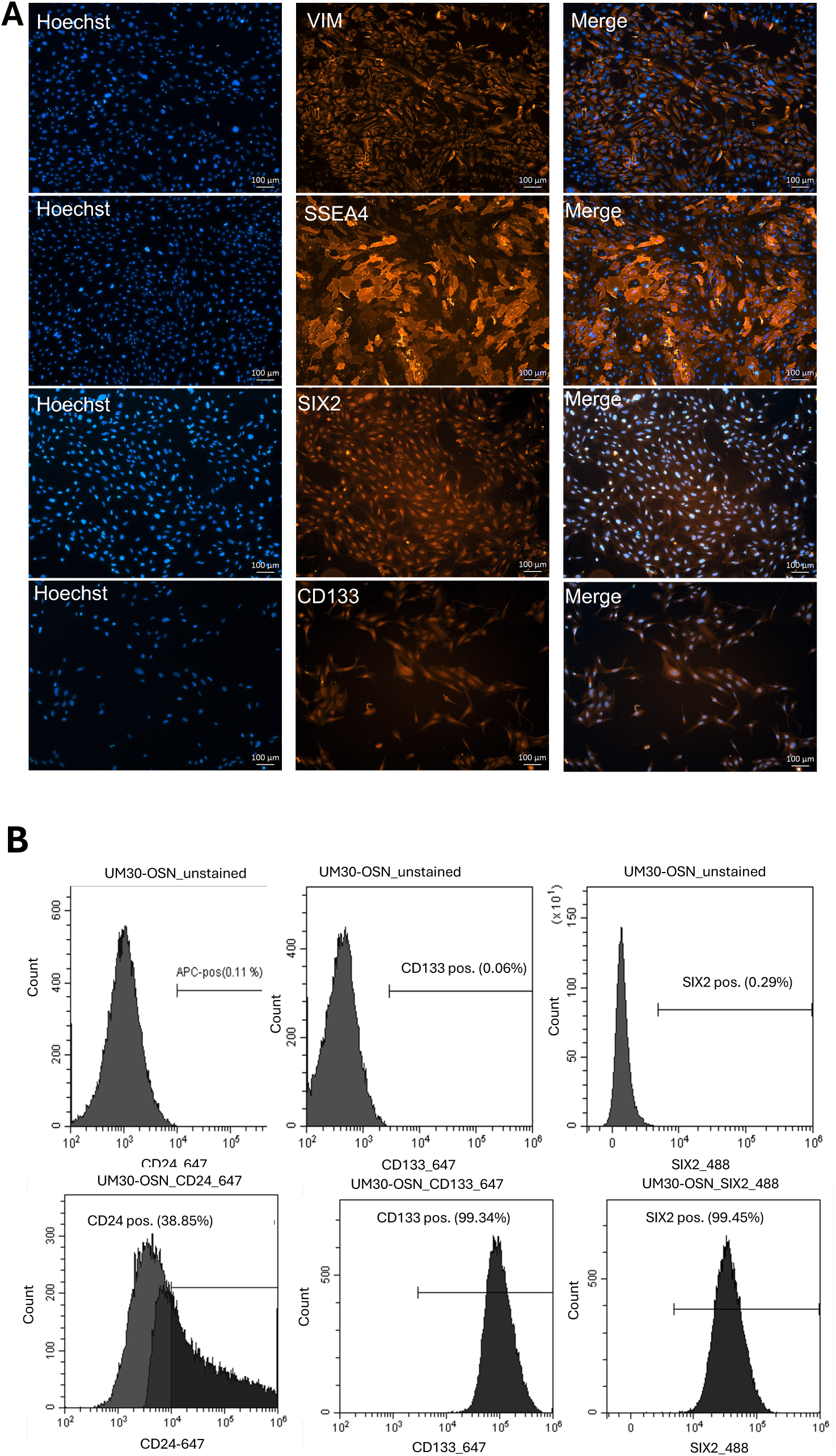
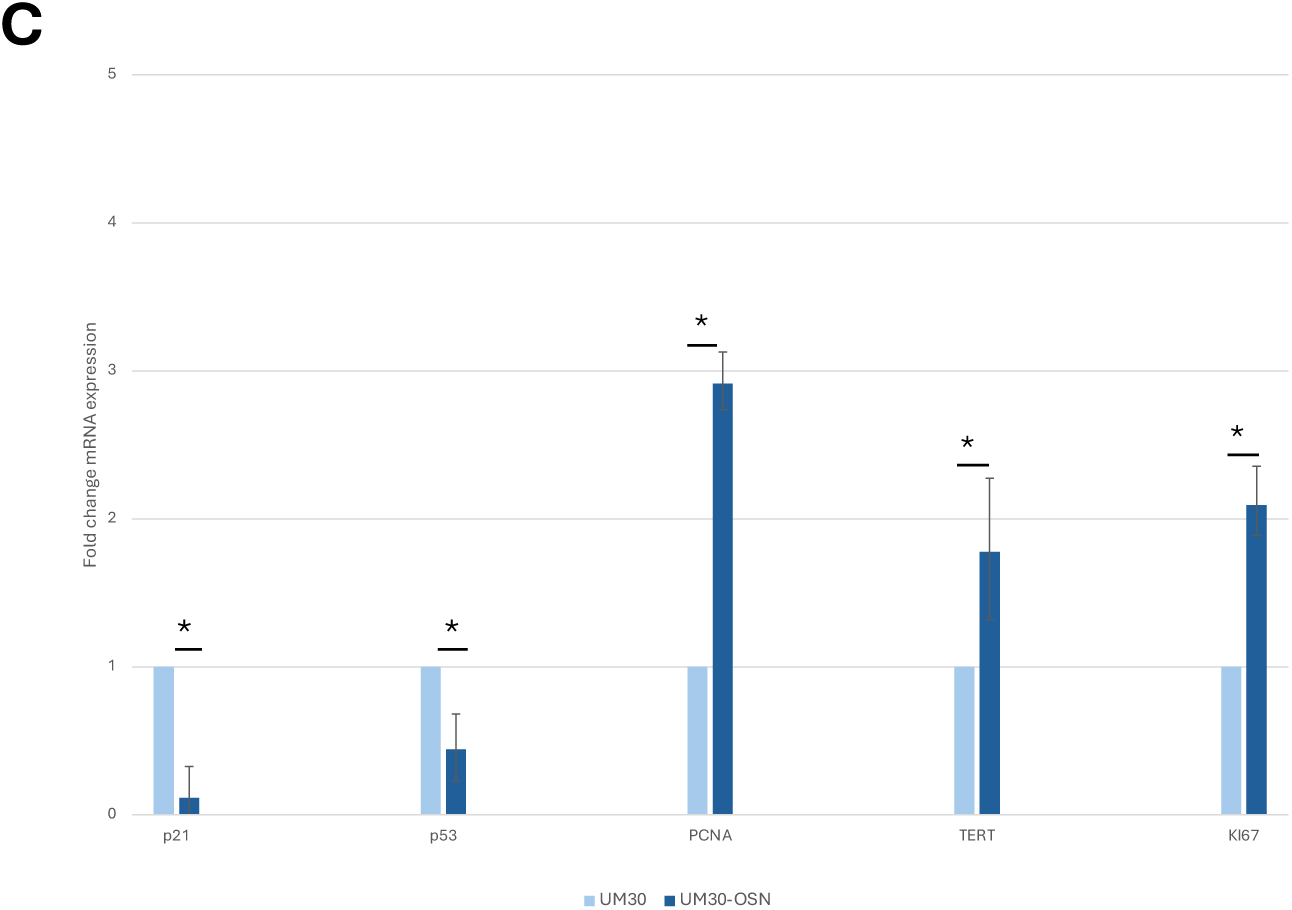
Characterization of immortalized UM30-OSN cell line. **(A)** The renal progenitor origin of the urine-derived UM30-OSN cell line was confirmed by immunocytochemistry for the renal stem cell markers SIX2 and CD133, the mesenchymal marker Vimentin, and the pluripotency marker SSEA4. **(B)** Flow cytometry (FACS) further validated the expression of SIX2, CD24, and CD133. **(C)** Quantitative real-time PCR was used to assess the expression of cell cycle-related genes, including p21, p53, hTERT, PCNA, and KI67. Gene expression levels were normalized to the housekeeping gene glycerinaldehyd-3-phosphat-Ddehydrogenase (GAPDH).

To further confirm stemness, fluorescence-activated cell analysis (FACS) was also performed (Figure 2B) and this revealed that approximately 99.45% of the cells express SIX2, CD133 (99.34%) and the renal progenitor cell surface marker CD24 (38.85%).

Immortalized cell lines are characterized by a high proliferation rate, which can be analyzed not only morphologically but also at the cellular level. Based on this, we comparatively analyzed the expression of the cell cycle-associated genes-*P53, P21, KI67, PCNA*, and *TERT* in UM30-OSN and UM30. UM30-OSN showed a significantly higher expression of *PCNA* (2.9-fold), *KI67* (2.0-fold), hTERT (1.77-fold), whilst the expression of P53 and P21 were downregulated by 0.44-fold and 0.11-fold respectively (Figure 2C).

### Differentiation potential of UM30-OSN into human podocytes

UM30-OSN were differentiated into podocytes following our previously published protocols ^28,29^ (Figure 3A). The human immortal podocyte (AB 8/13) cell line was used for comparison and benchmarking ^3^. Characterization and assessment of successful differentiation were demonstrated by immunocytochemistry, quantitative real-time PCR, and Western blotting (Figure 3B-D). Expression of podocyte markers NPHS1, NPHS2 and SYNPO were detected only upon podocyte differentiation of UM30-OSN.

**Figure 3:**
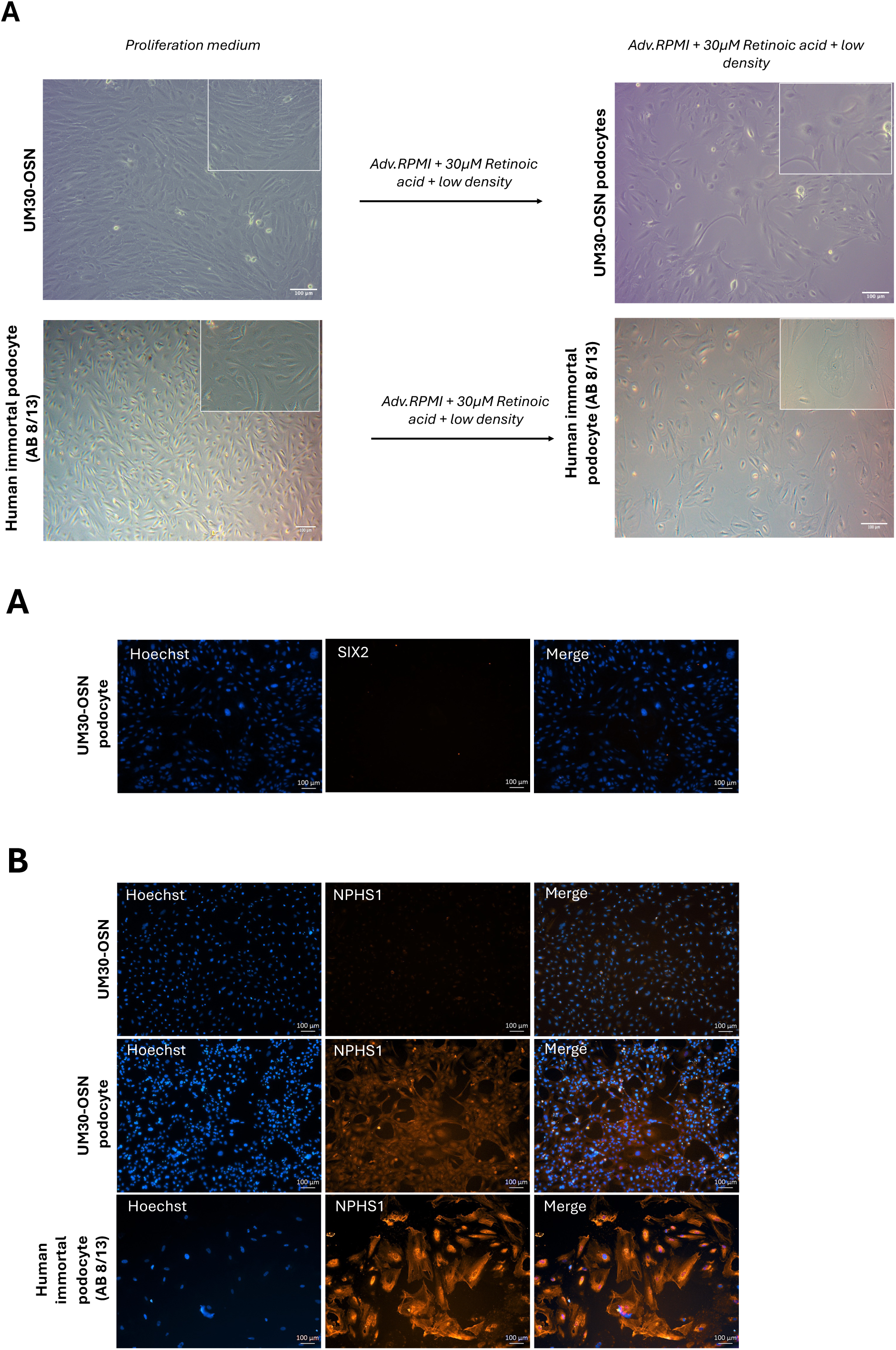
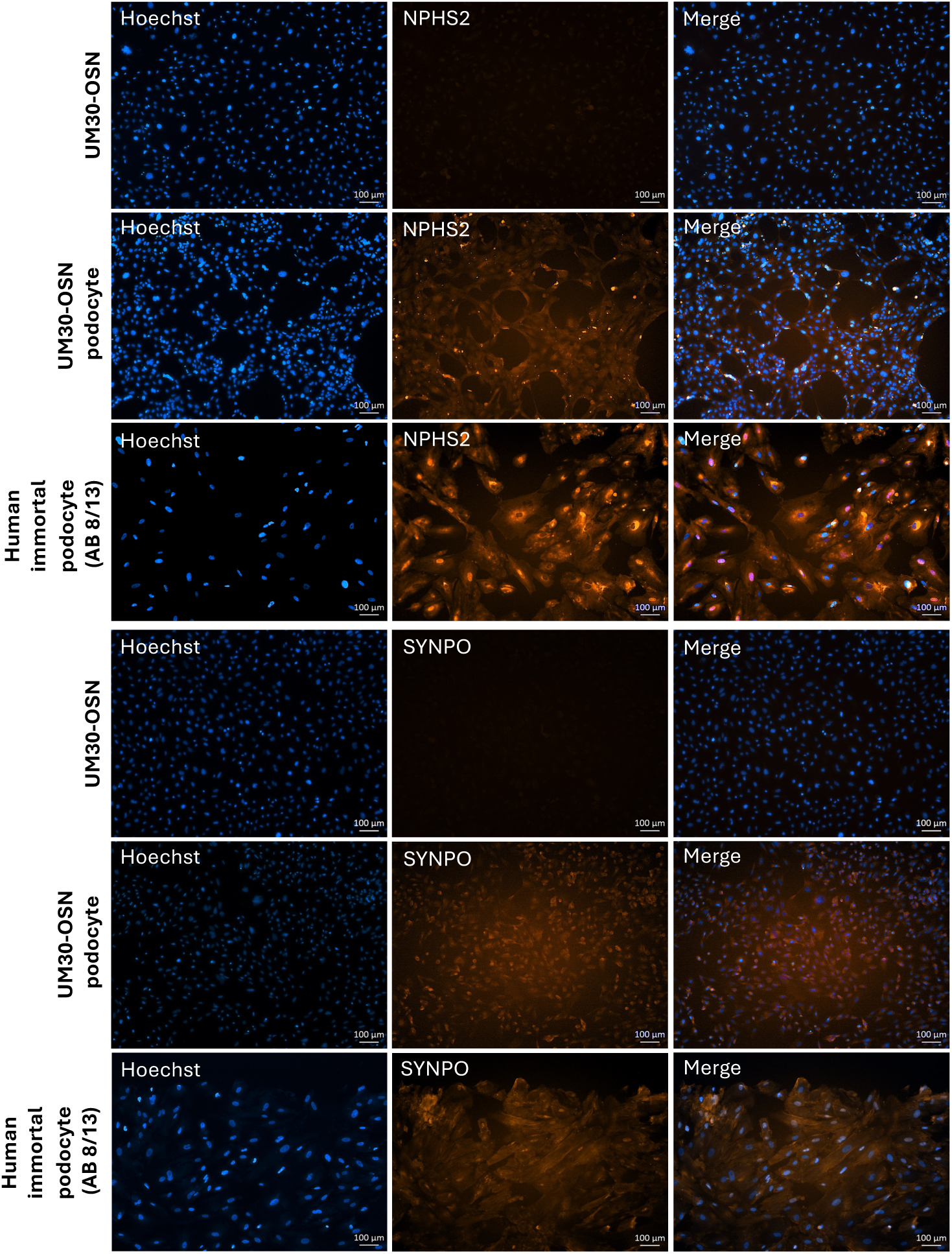
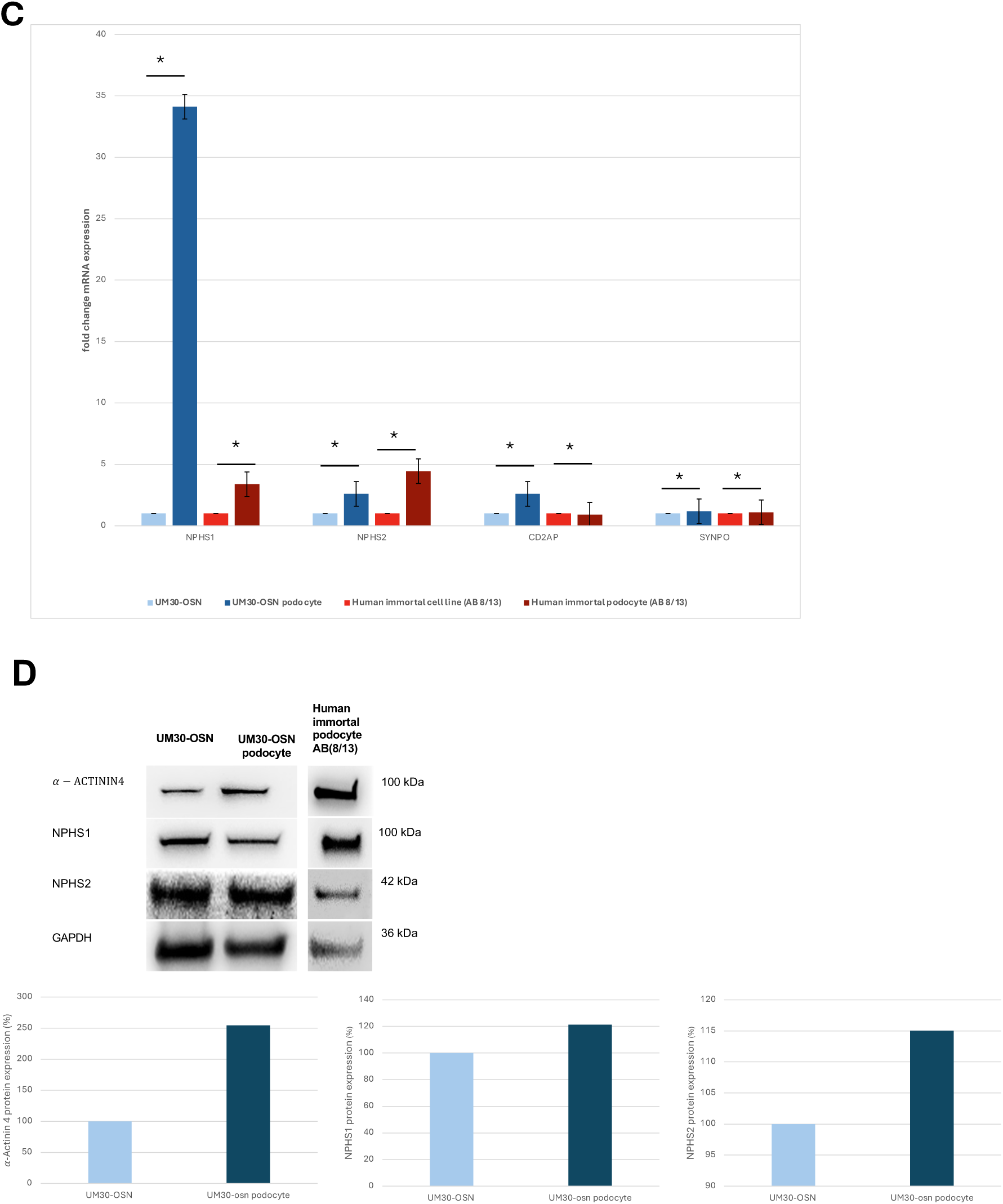
Culturing UM30-OSN Cells in advanced RPMI + RA induces podocyte differentiation. **(A)** UM30-OSN cells and human immortalized podocytes (AB8/13) were cultured in advanced RPMI medium supplemented with 30 µM retinoic acid (RA) to promote differentiation into mature podocytes. Light microscopy revealed the characteristic “fried egg” morphology. **(B)** Immunofluorescence staining confirmed the expression of podocyte-specific markers, including NPHS1, NPHS2 and α-ACTININ4 (scale bars: 100 μm). **(C)** Quantitative real-time PCR and **(D)** Western blot analysis further validated marker gene expression. GAPDH served as the internal control for normalization.

On the mRNA level, UM30-OSN podocytes show increased expression of the podocyte markers *NPHS1* (34.11 fold), *NPHS2* (4.19 fold), *CD2AP* (2.59 fold) and *SYNPO* (1.17 fold) compared to the undifferentiated UM30-OSN (Figure 3C).

Elevated mRNA expression of the podocyte markers *NPHS1* (3.37 fold), *NPHS2* (4.44 fold), *CD2AP* (0.90 fold) and *SYNPO* (1.10 fold) was also demonstrated in the immortal podocytes (AB 8/13) cultured at 37C compared cells cultured at 33C (Figure 3C). These results corroborate the protein-based analyses (Figure 3D). The housekeeping protein Glyceraldehyde −3-phosphate-dehydrogenase (GAPDH) served as loading control.

### Comparative transcriptome and Gene Ontology analysis of podocytes differentiated from UM30, UM30-OSN and the immortal AB 8/13 cell line

Hierarchical cluster analysis was carried out to compare similarities between their transcriptomes (Figure 4A, 4B). The correlation table shows a higher correlation co-efficient between UM30 and UM30-OSN podocytes as expected (R^2^ = 0.94). However, this value drops to 0.88 when comparing UM30 and AB 8/13 podocytes (Figure 4B). Additionally, a Venn diagram-based transcriptome comparison was carried out between UM30-OSN and AB 8/13 podocytes (Figure 4C). A total of 14947 genes were detected as expressed in common expressed in both podocyte lines (Figure 4C). Comparing the expressed genes (det-p < 0.05), 1341 genes are exclusively expressed in AB 8/13 podocytes and 2249 genes in UM30-OSN podocytes (Figure 4C).

**Figure 4:**
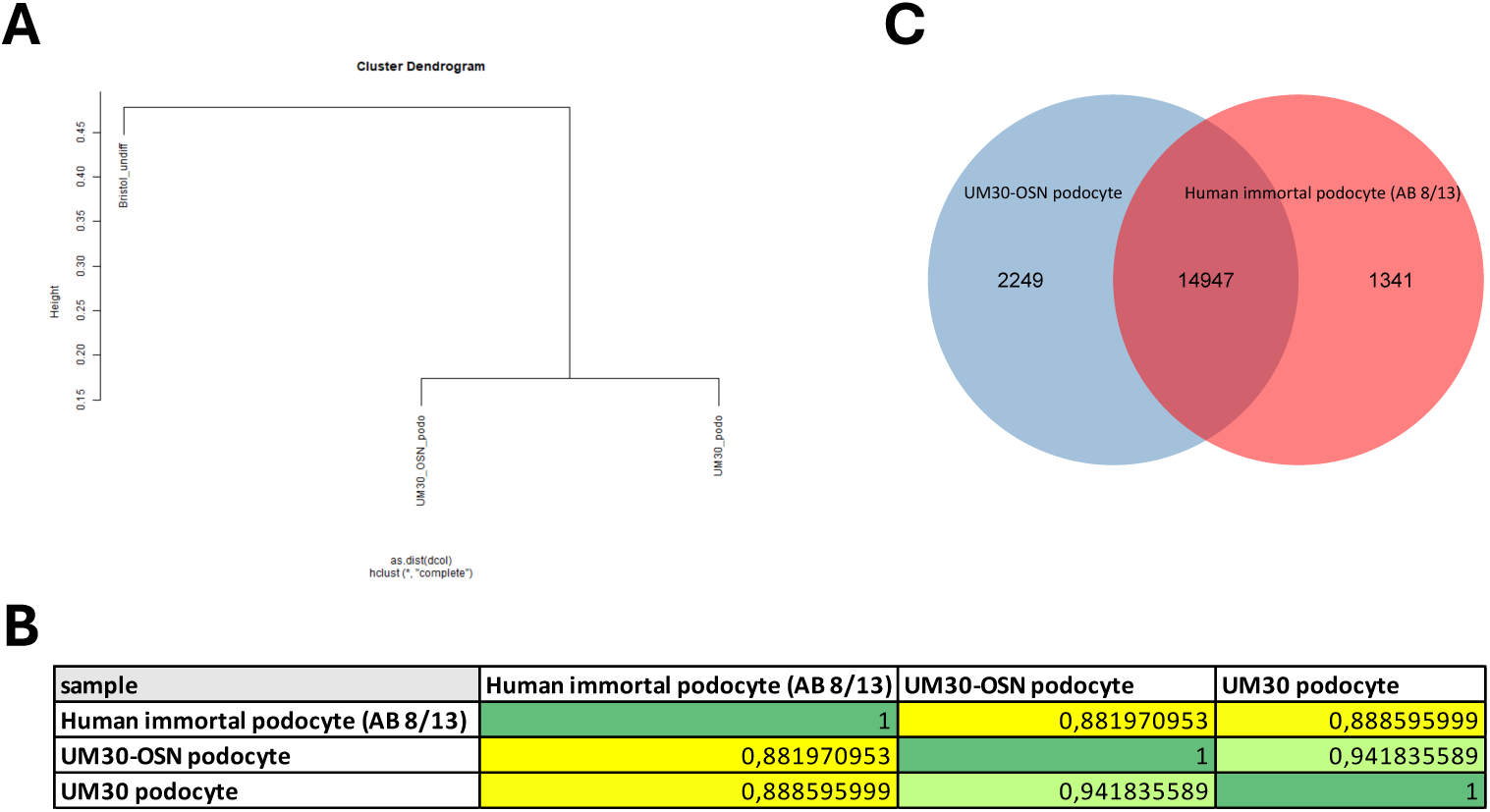
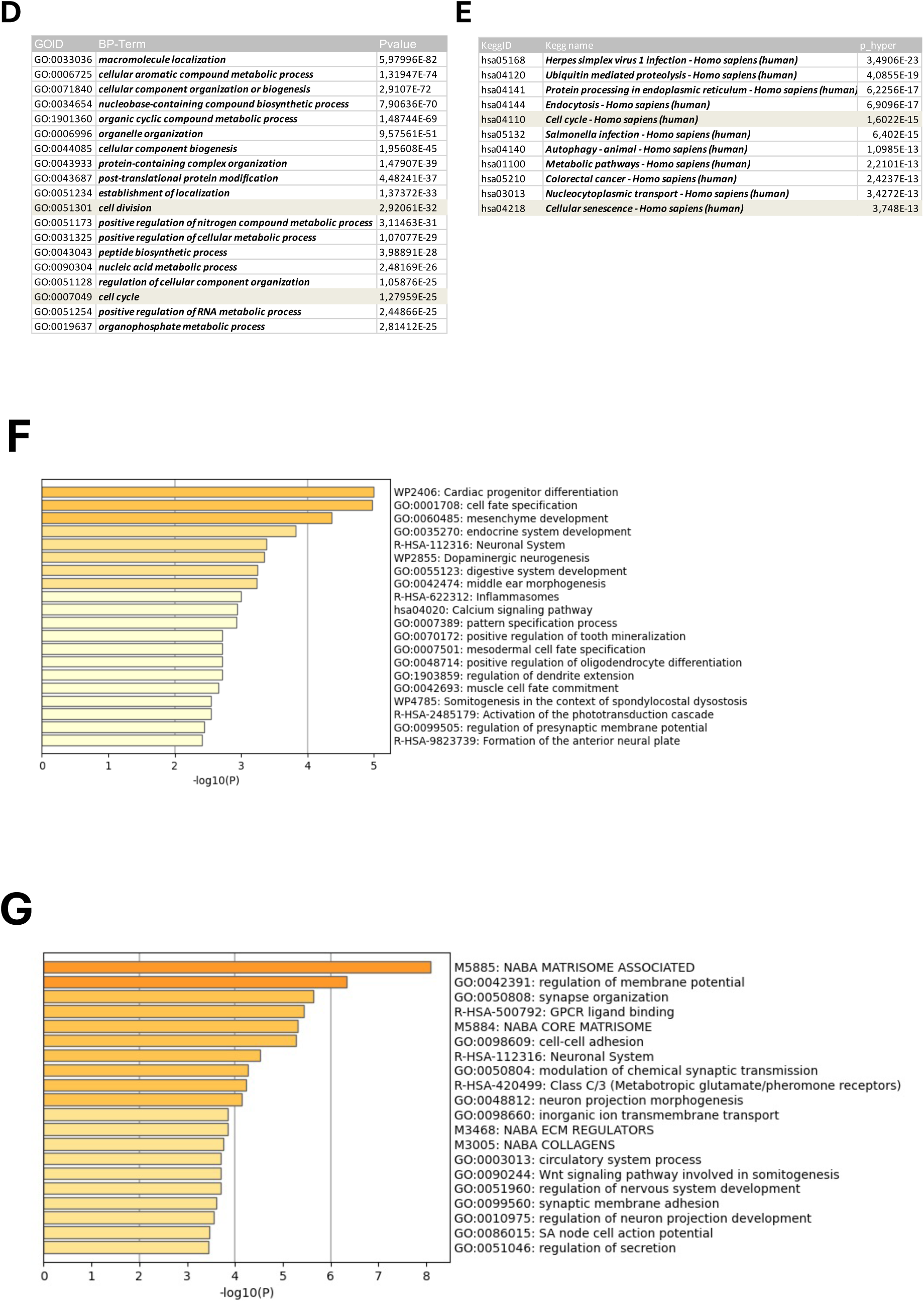
Comparative transcriptome and Gene Ontology analysis of human podocyte cell lines. **(A)** A cluster diagram between the human immortal podocytes (AB 8/13), urine-derived UM30 podocytes and the UM30-OSN-derived podocytes was performed. **(B)** The correlation table between the UM30-OSN and the immortal podocytes (AB 8/13) revealed a correlation coefficient of 0.88. **(C)** A Venn diagram was performed and shows 14947 common genes expressed in common between the two cell types**. (D-E)** The common GO BP-terms and KEGG pathways between the immortal podocytes (AB 8/13) and UM30-OSN podocytes are associated with cell cycle, cell division and cellular senescence. **(F)** Metascape analysis of the 1341 genes only expressed in the immortal podocytes (AB 8/13). **(G)** Shows the exclusively 2249 genes expressed only in the UM30-OSN podocytes by metascape analysis.

The common GO:BP terms and pathways related to the 14947 genes expressed in both podocyte cell lines are associated with cell division, cell cycle and cellular senescence (Figure 4D-E). The entire GO BP-terms and KEGG pathway is presented in Supplemental Table S3 and S5. Metascape analysis related to the 1341 genes exclusively expressed in the human immortal pododcyte line (AB 8/13) are e.g. mesenchymal development, calcium signaling and neuronal system. In parallel the exclusive GO:BP terms and pathways of the UM30-OSN podocytes related to the 2249 genes are e.g. regulation of membrane potential, synapse organization, cell-cell-adhesion and synaptic membrane adhesion. The full gene list can be found in Supplementary Table S4B.

### Modulation of APOL Gene Expression and Kidney Injury Markers by IFN-γ and Baricitinib in Human UM30-OSN Podocytes

Upon IFN-γ binding, the canonical signaling cascade is initiated via the JAK/STAT pathway, predominantly through activation of JAK1 and JAK2, which are linked to the intracellular domains of the IFN-γ receptor (Figure 5A). To investigate the effect of IFN-γ on the UM30-OSN podocytes, the cells were pre-treated with or without the JAK1/2 inhibitor -Baricitinib upon IFN-γ stimulation (Figure 5B). Baricitinib inhibited IFN-γ-induced morphological and molecular alterations in UM30-OSN podocytes. Treatment of UM30-OSN podocytes with 100 ng/ml IFN-γ for 24 hours led to a pronounced increase in rounded, detached cells compared to the untreated control, as observed by light microscopy. Co-treatment with Baricitinib significantly reduced the number of detached cells, resulting in a morphology comparable to that of control cells (Figure 5C). To assess the molecular response to IFN-γ and its modulation by Baricitinib, we analyzed the expression of IFN-γ-responsive genes-APOL1, MAPK8, STAT3 and kidney injury markers Cycs, IL-6, NGAL, CLU, Caspase-1, TGF-β and Gasdermin D using quantitative real-time PCR (Figure 5D). IFN-γ treatment significantly upregulated several of these genes including APOL1, whereas co-treatment with Baricitinib largely attenuated this induction, restoring expression levels closer to those of the control. Western blot analysis confirmed the transcriptional findings at the protein level. IFN-γ induced phosphorylation of STAT1 (p-STAT1) thus leading to activated expression of APOL1 and down-regulation of the podocyte marker-α-ACTININ4. Baricitinib co-treatment reversed these effects, preserving α-ACTININ4 expression and reducing APOL1 and p-STAT1 to near-baseline levels (Figure 5E). Immunofluorescence staining for fibrosis-associated proteins Fibronectin, α-SMA, and Vimentin further supported the deleterious effects of IFN-γ. All three markers were upregulated following IFN-γ treatment, thus indicating a shift towards a pro-fibrotic phenotype. This effect was absent in control and Baricitinib-treated cells, suggesting a protective role of Baricitinib against IFN-γ-induced fibrosis (Figure 5F).

**Figure 5:**
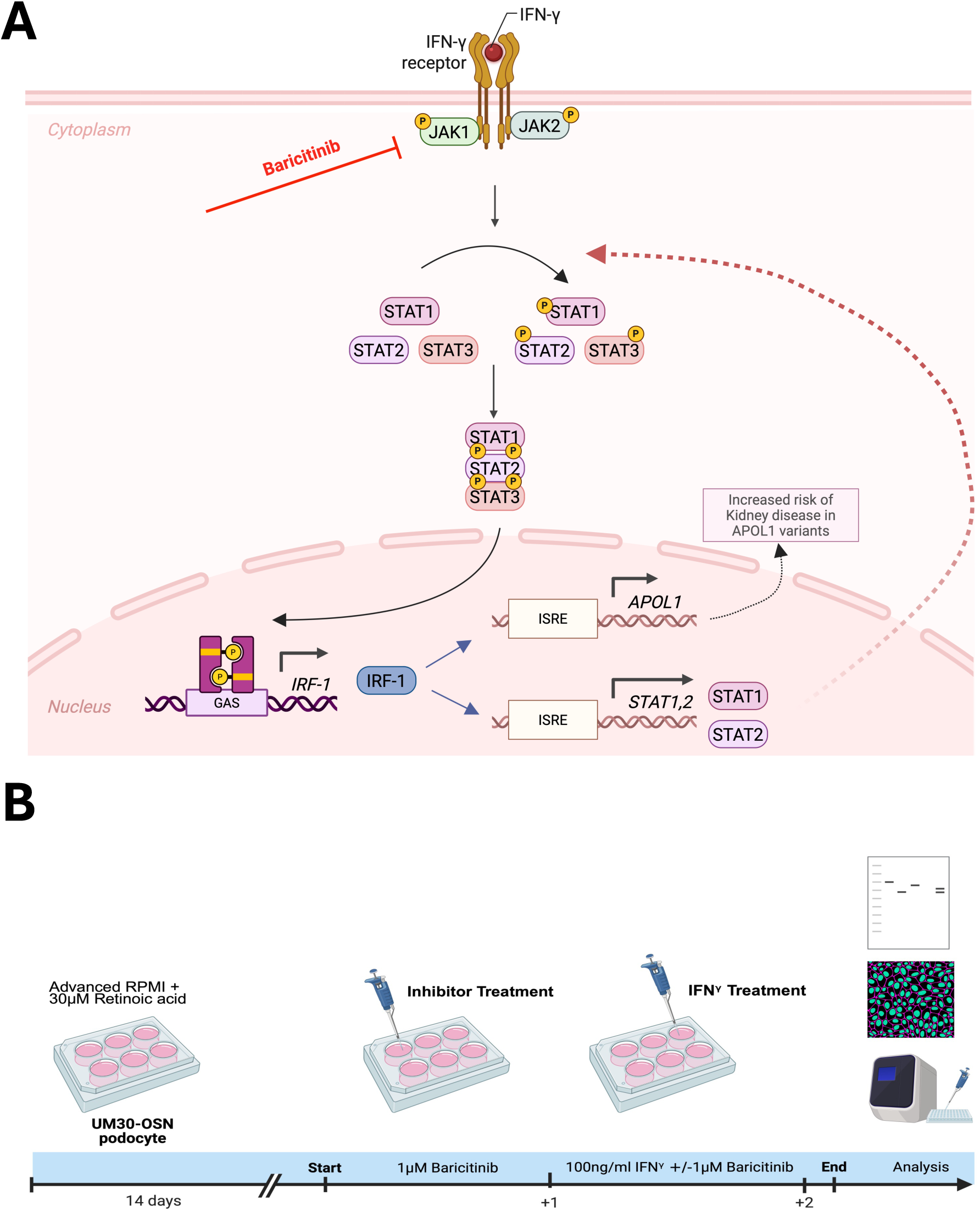
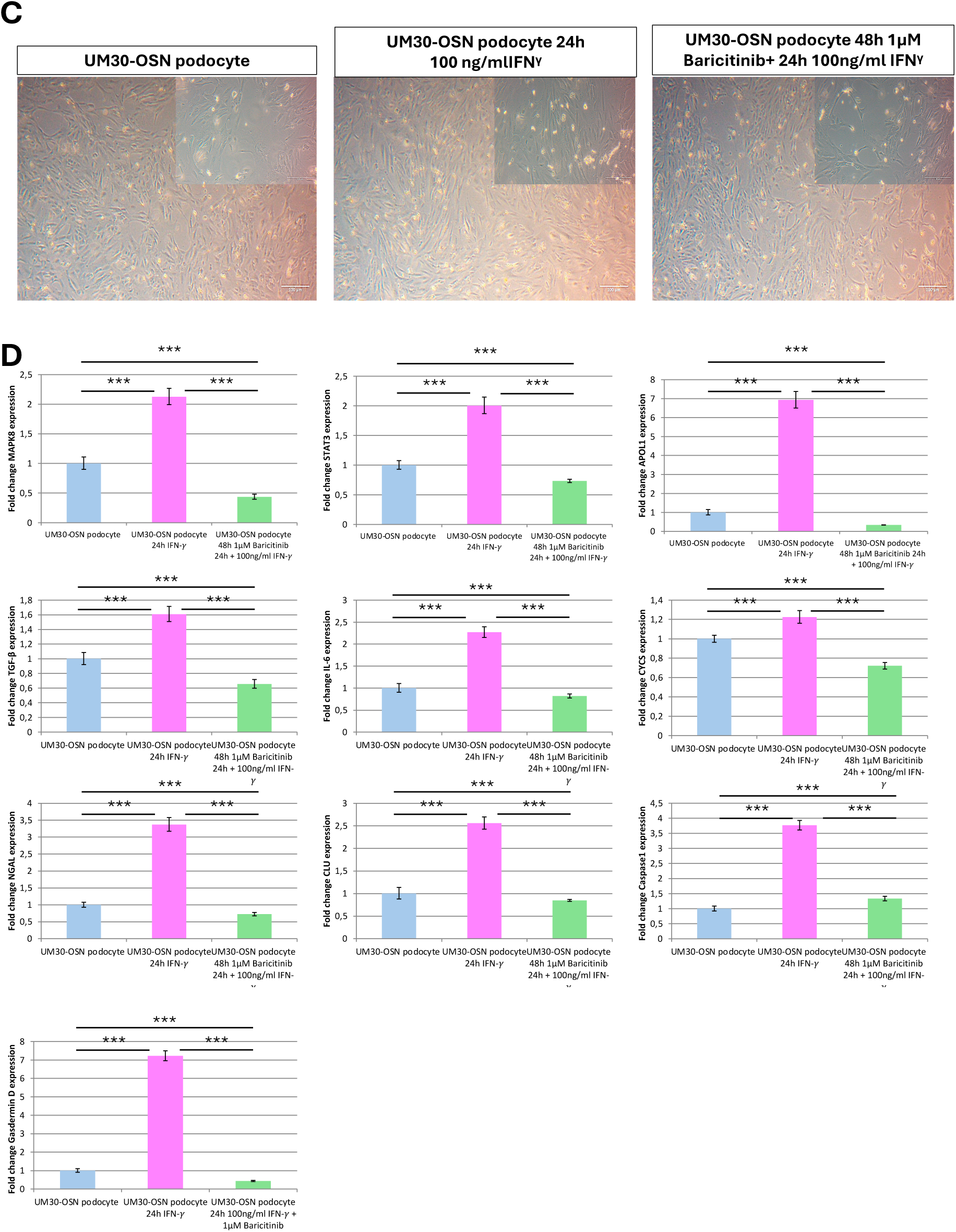
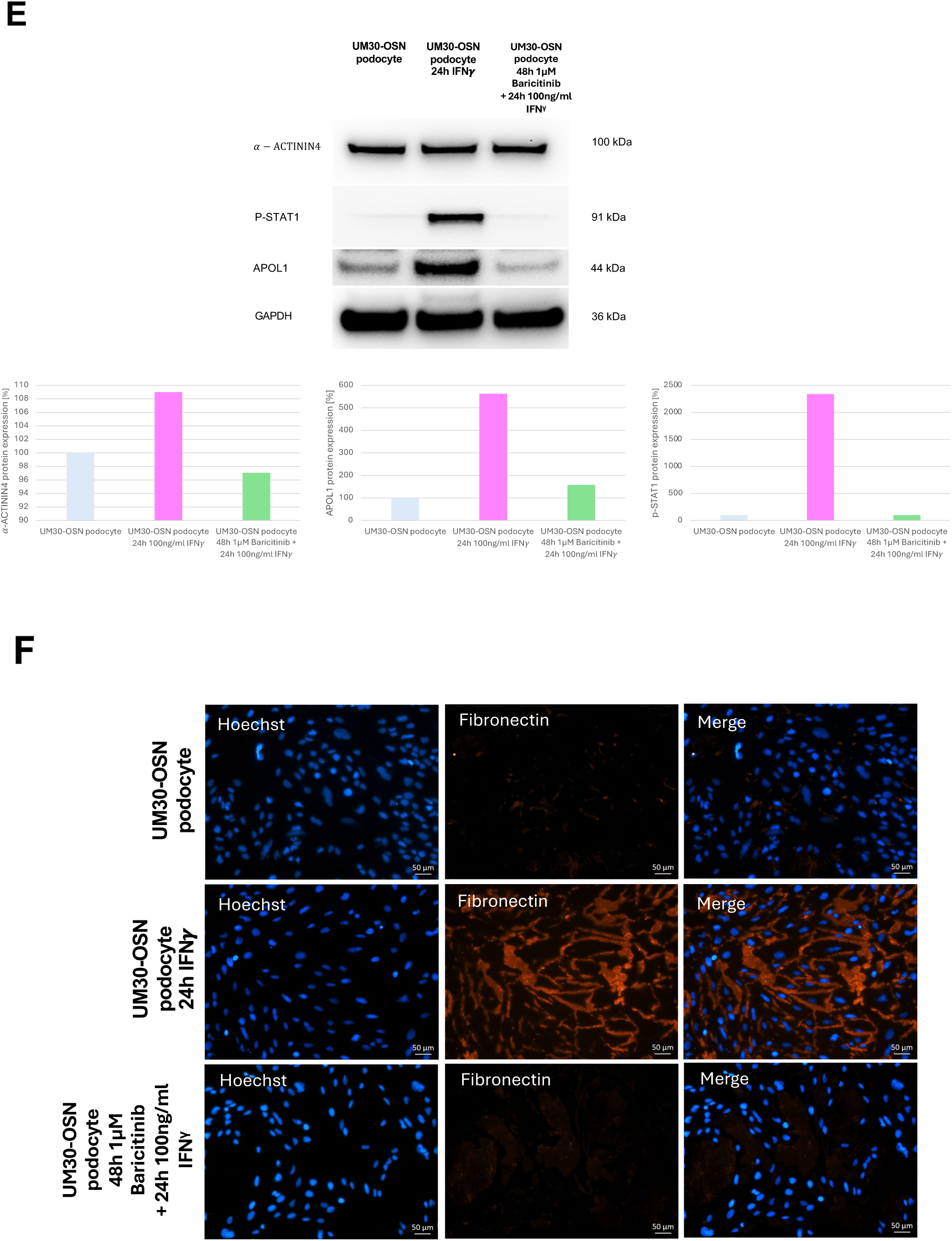
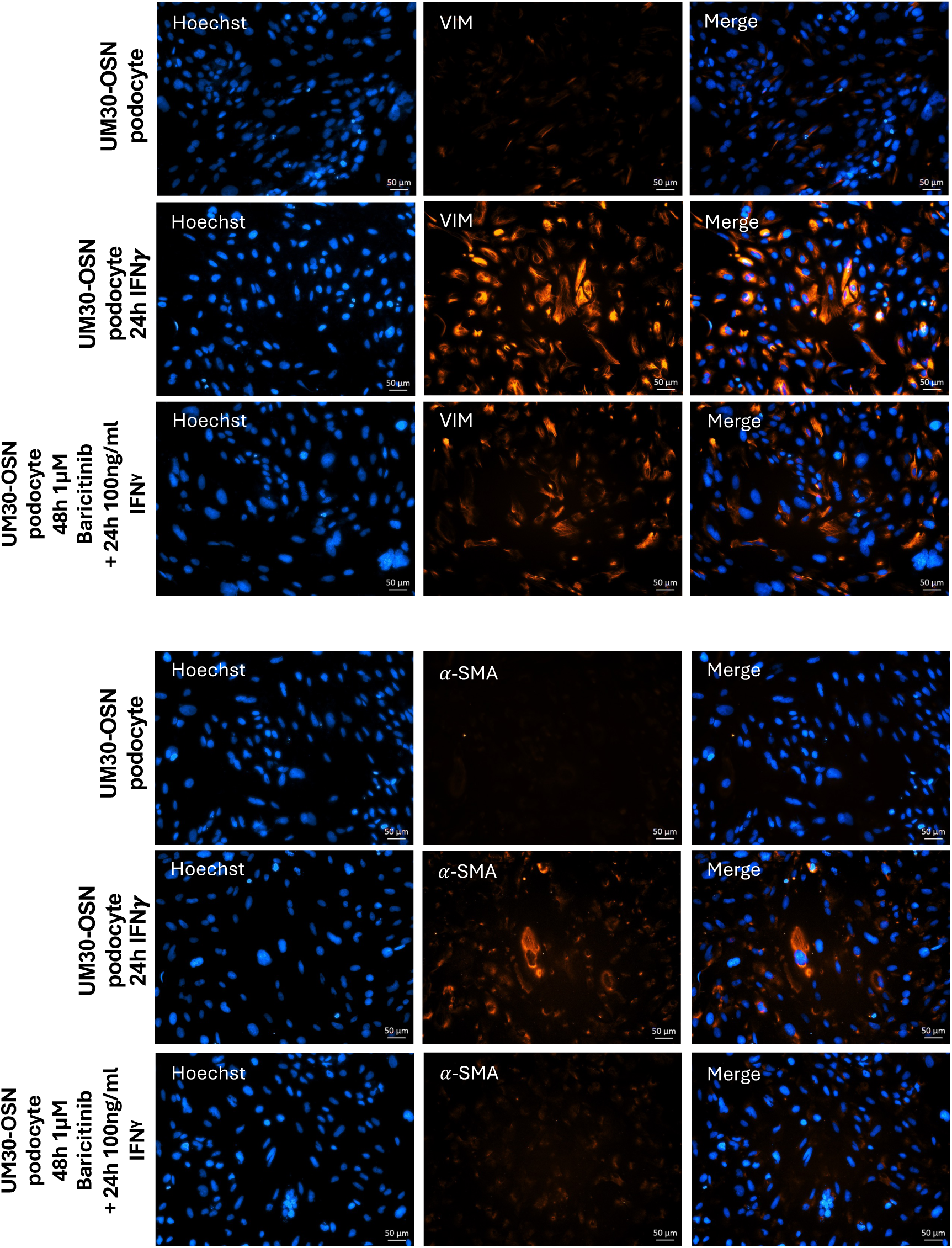
Effects of IFNγ and Baricitinib on UM30-OSN podocytes APOL gene expression and kidney injury marker. (A) Schematic representation of the IFN-γ signaling pathway. Binding of interferon-gamma (IFN-γ) to its heterodimeric receptor, composed of IFNGR1 and IFNGR2, induces receptor dimerization and subsequent activation of the associated Janus kinases JAK1 and JAK2 via transphosphorylation. This leads to the recruitment and phosphorylation of the signal transducers STAT1, STAT2, and STAT3. Phosphorylated STATs dimerize and translocate into the nucleus, where they bind specific DNA elements such as GAS or ISRE sequences and activate the transcription of target genes (e.g., IRF-1, STAT1/2, APOL1). Baricitinib inhibits upstream signaling by selectively blocking JAK1/JAK2 activity, thereby suppressing STAT activation and downstream gene expression. (B) Experimental setup using immortalized urine-derived renal progenitor cells differentiated into podocytes (UM30-OSN). After 14 days of podocyte differentiation, cells were pre-treated with or without 1 μM Baricitinib for 24 h, followed by stimulation with 100 ng/ml IFN-γ for an additional 24 h. Subsequent analyses were performed on both RNA and protein levels. (C) A 24-hour treatment with 100 ng/ml IFN-γ resulted in an increased number of rounded (detached) cells compared to untreated controls, as visualized by light microscopy. The combination of IFN-γ with Baricitinib significantly reduced the number of rounded cells, resembling the control condition. (D) The expression of IFN-γ-associated genes such as APOL1, MAPK8, STAT3, and kidney injury markers such as Cycs, IL-6, NGAL, CLU, Caspase-1, TGF-β, and Gasdermin D was analyzed by quantitative real-time PCR under all three conditions. (E) Western blot analysis was performed to assess protein levels of the podocyte marker alpha-ACTININ4, APOL1, and phosphorylated STAT1 (p-STAT1). (F) Immunofluorescence-based detection of Fibronectin, alpha-SMA, and Vimentin indicated fibrosis-like tendencies upon IFN-γ treatment, evidenced by the upregulated expression of all three proteins. No such upregulation was observed in control or Baricitinib co-treated cells (Scale bars: 100 μm). Figure 5A-B were created in https://BioRender.com.

**Figure 6:**
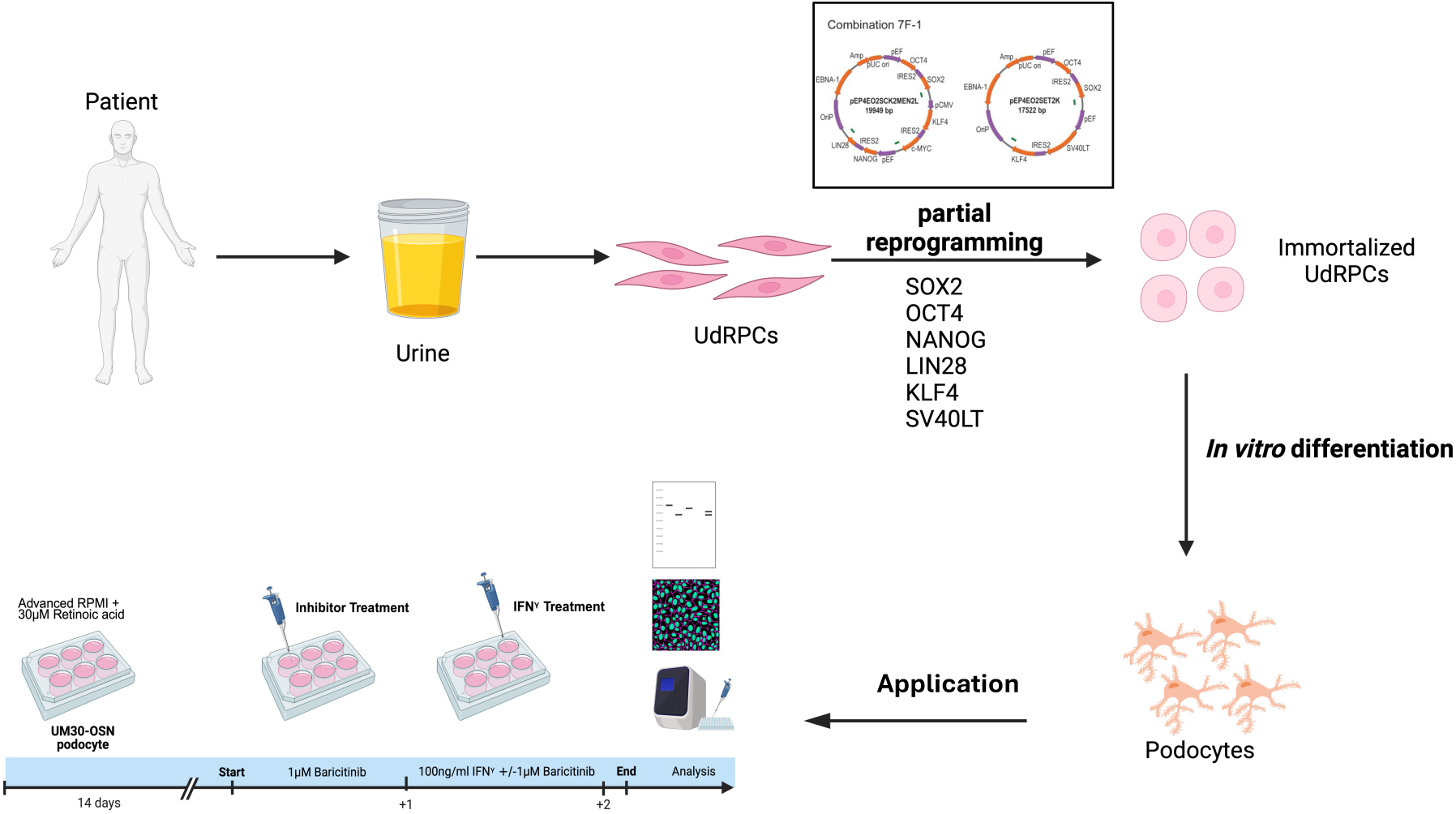
Generation of Immortalized SIX2⁺ Renal progenitor Cells from urine of a West African male. SIX2-positive urine-derived renal progenitor cells (UdRPCs) from a 30 years old male of west African origin were isolated from his urine samples and subjected to episomal-based cellular reprogramming. Although full pluripotency was not induced, the cells acquired a significantly enhanced proliferative capacity compared to primary UdRPCs. Notably, this immortalized cell population retains the ability to differentiate into functional human podocytes, offering a robust and scalable model for renal research. For further application, the human podocytes were treated with and without 1µM Baricitinib, followed by a 24h stimulation with 100ng/ml IFNγ. UM30-OSN represents and a disease-relevant model for studying APOL1-mediated kidney fibrosis. The figure was created using BioRender (https://BioRender.com.)

## Discussion

As mature podocytes are terminally differentiated cells, it is not possible for them to undergo cell division and therefore have a limited life-span in culture^46^. To circumvent this shortfall, we previously established protocols for isolating highly proliferative SIX2-positive urine derived progenitor cells ^25^ and others have also isolated urine-derived epithelial cells ^47^. As these cells also have are endowed limited life-span in culture, this has been by-passed by immortalizations for use in further studies related to nephrogenesis and modelling kidney-associated diseases ^1–3,47–49^. The available cell lines have so far been derived from individuals of Caucasian heritage, except the UM51 hTERT ^2^.

In the present manuscript, we present the successful immortalization of a human SIX2-positive renal progenitor cell line derived from the urine of a 30-year-old West African male (UM30-OSN). These cells show stable morphology and proliferation for over one year in culture (∼100 passages). To achieve partial reprogramming and immortalization, plasmids encoding SV40 and the reprogramming factors-OCT4, SOX2, NANOG, cMyc, and KLF4 were employed.

The conferral of unlimited cell division is associated with chromosomal alterations and genomic instabilities ^50,51^. In the UM30-OSN these are also present in some cases, whereby P53, P21 as well as PCNA and KI67 expression and their activation and downregulation indicate intact cell cycle checkpoints. The comparison of the transcriptomes of UM30, UM30-OSN and the UM51 iPSC showed a clear separation, in which UM30-OSN clustered closer to the primary UdRPCs UM30 than to the iPSC.

UM30-OSN retains its renal origin due to partial reprogramming. This is confirmed by the expression of vimentin and renal progenitor stem cell, SIX2 and cell surface markers CD24, CD133 [36]. The lack of expression of the pluripotency markers-POU5F1, NANOG, SOX2 and CDH1 further confirms that UM30-OSN is not pluripotent.

Shared KEGG pathways and GO-Biological Process (GO-BP) terms identified in both immortalized AB8/13 podocytes and UM30-OSN podocytes are associated with cell division, cell cycle regulation, and cellular senescence. These biological processes are well-established hallmarks of cellular immortalization, thereby supporting the molecular basis of the immortalization observed in these cell lines ^52^. Differentiation into mature podocytes was shown based on protein and mRNA expression of Nephrin (NPHS1), Podocin (NPHS2), Synaptopodin (SYNPO) and CD2-associated protein (CD2AP). According to the present findings, numerous podocyte-specific genes have been identified whose products play an important role in maintaining the filtration barrier. Near to the extracellular proteins of the slit membrane is the hairpin-like transmembrane protein NPHS2 with two cytoplasmic ends, which is expressed within the glomerulus only in podocytes ^53^. An interaction between NPHS2, NPHS1 and CD2AP has already been demonstrated ^54,55^. This highly conserved complex, through which podocytes are connected to cell-cell contacts, controls podocyte function ^56,57^. Besides the essential components of the podocyctic slit membrane, the components the podocytic cytoskeleton and associated molecules are necessary for a mature podocyte. SYNPO a protein that interacts with several cytoskeletal and membrane-associated molecules is restricted expressed in differentiated podocytes in the kidney ^58^. It has relevance in the maintenance of podocyte morphology.

Thus, the expression of podocyte-specific genes, by culturing UM30-OSN in adv. RPMI and RA, confirm the differentiation potential of this cell line into podocytes. In summary, we have successfully described the first male African cell line UM30-OSN immortalized with SV40 and the reprogramming factors OCT4, SOX2, NANOG, cMyc and KLF4. These grow as monolayers and have a rice grain-like morphology. and can be differentiated into mature podocytes amenable for studying nephrogenesis and its kidney-associated diseases.Thus, in addition to iPSC and the human immortal (AB 8/13) podocytes, they serve as an additional cell culture option with a major advantage of West African origin.

Beyond highlighting UM30-OSN cells as a viable alternative to commonly used cell lines of Caucasian origin such as HEK and the immortalized podocyte cells, our study also supports their application as a disease-relevant *in vitro* model for investigating APOL1-associated nephropathies.

In this study, we demonstrate that Baricitinib effectively counteracts the harmful effects of IFN-γ on human urine-derived UM30-OSN podocytes, both at the molecular and morphological levels. Treatment with IFN-γ led to marked changes in cell morphology, including detachment and rounding, along with increased expression of APOL1 and other markers associated with cellular stress, inflammation, and fibrosis. Our findings are consistent with previous reports describing IFN-γ as a potent inducer of APOL1 expression via activation of the JAK/STAT signaling pathway ^11,16,59–62^.

Specifically, we demonstrated that IFN-γ induces a characteristic rounding and detachment of podocytes, accompanied by upregulation of fibrosis markers such as Fibronectin, α-SMA, and Vimentin, as well as APOL1 and additional stress- and inflammation-associated genes, such as STAT3, IL-6, MAPK8, Caspase-1, and Gasdermin D. In particular Caspase-1 and Gasdermin D are associated with cellular immune response and inflammatory reactions and are key players in the pyroptosis signaling pathway, a form of programmed cell death ^63,64^.

These findings are similar with previously reported mechanisms by which IFN-γ promotes a pro-fibrotic and cytotoxic environment mediated through activation of the JAK/STAT signaling pathway.^11,65^.

Baricitinib effectively prevented these IFN-γ-induced effects. Podocyte morphological integrity was preserved, APOL1 expression and STAT1 phosphorylation were reduced, and fibrosis markers remained at baseline levels. This protective effect is consistent with observations in human kidney biopsies and podocyte models, where Baricitinib has been shown to reduce cytotoxicity and APOL1 expression under pro-inflammatory conditions ^10^. Mechanistically, our Western blot results showing reduced p-STAT1 expression confirm effective blockade of the JAK/STAT phosphorylation cascade, which plays a central role in APOL1 transcription and downstream cellular injury. These findings highlight the therapeutic potential of Baricitinib in APOL1-associated kidney diseases, as applied in the Phase 2 clinical JUSTICE trial, which evaluated its efficacy in reducing proteinuria in individuals carrying high-risk APOL1 genotypes ^60^.

## Conclusion

In conclusion, our results confirm that IFN-γ induces a pro-fibrotic and APOL1-driven stress response in UM30-OSN derived podocytes via JAK/STAT signaling. Baricitinib acts as an effective inhibitor of this signaling cascade, highlighting its strong potential as a therapeutic option for APOL1-associated kidney diseases. Ultimately, this study not only underscores the utility of the urine-derived immortalized UM30-OSN cell line, but also establishes its relevance as a valuable *in vitro* model for investigating APOL1-mediated kidney diseases such as fibrosis.

## Supporting information

Supplemtal Material Table of Contents, Supplementary Table S3, S4, S5

## Declarations

## Author Contributions

Conceptualization, J.A.; methodology, C.T., R.M.; bioinformatics, W.W.; formal analysis, C.T.; investigation, C.T., W.W.; resources, J.A., O.A.; data curation, C.T.; writing—original draft preparation, C.T.; writing—review and editing, C.T. and J.A.; visualization, C.T.; supervision, J.A.; project administration, J.A; funding acquisition, J.A. All authors have read and agreed to the published version of the manuscript.

## Institutional Review Board Statement

The study was conducted in accordance with the written approval (Ethical approval Number: 2017-2457_3) of the ethical review board of the medical faculty of Heinrich Heine University, Düsseldorf, Germany. All methods were carried out in accordance with the approved guidelines. Medical faculty of Heinrich Heine University approved all experimental protocols.

## Informed Consent Statement

In this study, urine samples were collected with the informed consent of the donors.

## Data Availability Statement

Gene expression data will be available online in the National Centre of Biotechnology Information (NCBI) Gene Expression Omnibus under the number GSE299018.

## Acknowledgments

James Adjaye acknowledges funding from the medical faculty of Heinrich Heine University, Duesseldorf, Germany. Dr. Ania Koziell and Professor Saleem Moin provided the human immortal cell line (AB 8/13). Dr. Nina Graffmann assisted with the FACs analyses. Martina Bohndorf assisted by initiating the partial reprogramming experiments.

