## Supplementary material for "Modelling APOL1-mediated kidney inflammation and fibrosis using a partially reprogrammed urine derived SIX2-positive renal progenitor cell line": Supplemtal Material Table of Contents, Supplementary Table S3, S4, S5: Supplemental Material Table of Contents.docx

**Supplementary Materials**

**Supplementary Table S1: Used antibodies.**

**Supplementary Table S2: Used qPCR primers.**

**Supplementary Table S3: gostats.**

**Supplementary Table S4: full gene list. (a) full gene list corresponding to the Venn diagram presented in Figure 4C, showing the differences of the UM30-OSN and human immortal podocyte (AB 8/13) and their similarities. (b) full gene list corresponding to Figure 1H showing the differences between UM30, UM30-OSN and the UM51 iPSC and their similarities.**

**Supplementary Table S5: KEGG signaling pathways.**

**Supplementary Figure S1: uncropped western blot images.**

**Supplementary Figure S2: Plasmid maps.**

**Supplementary Figure S3: Karyogramm of UdRPC UM30-OSN and immortal cell line (AB 8/13).**

**Supplementary Table S1: Used antibodies**

| **Primary Antibody** | **Specificity** | **Dilution**  **IF FACS WB** | **Manufacturer** |
| --- | --- | --- | --- |
| **OCT4** | **Rabbit** | **1:200** | **Cell Signaling Technologies #2840S** |
| **SOX2** | **Rabbit** | **1:200** | **Cell Signaling Technologies #3579S** |
| **NANOG** | **Rabbit** | **1:200** | **Cell Signaling Technologies #4903S** |
| **Vimentin** | **Rabbit** | **1:200** | **CST**  **#5741** |
| **SSEA4** | **Mouse** | **1:200** | **Cell Signaling #MC813** |
| **SIX2** | **Mouse** | **1:200** | **Abnova**  **#H0010756-M01** |
| **CD133** | **Rabbit** | **1:200 1:50** | **Invitrogen**  **#PA5-38014** |
| **CD24** | **Mouse** | **1:50** | **Invitrogen**  **#MA5-11828** |
| **NPHS1** | **Rabbit** | **1:200** | **Invitrogen**  **#PA5-20330** |
| **NPHS2** | **Mouse** | **1:200** | **Sigma**  **#SAB4200810** |
| **SYNPO** | **Rabbit** | **1:200** | **Invitrogen**  **#PA-15794891** |
| **GAPDH** | **Mouse** | **1:1000** | **Ambion**  **#AM4300** |
| **HOECHST** |  | **1:5000** | **Thermo Fisher Scientific**  **#H3569** |
| **𝛼-SMA** | **Mouse** | **1:200** | **Dako**  **#0851** |
| **Fibronectin** | **Mouse** | **1:200** | **BD Transduction Laboratories**  **#BD610078** |
| **Vimentin** | **Rabbit** | **1:200** | **CST**  **#5741S** |
| **APOL1** | **Mouse** | **1:200 1:1000** | **Proteintech**  **#66124-1-iG** |
| **p-STAT1** | **Rabbit** | **1:1000** | **Cell Signaling**  **#9167S** |
| **p-JAK1/JAK2** | **Mouse** | **1:1000** | **Cell Signaling**  **#66245S** |
| **Secondary Antibody** |  |  |  |
| **Alexa 488** | **Rabbit** | **1:500** | **Thermo Fisher Scientific # A2106** |
| **Alexa 488** | **Mouse** | **1:500** | **Thermo Fisher Scientific #A11029** |
| **Goat anti-Rabbit IgG secondary antibody HRP** | **Rabbit** | **1:1000** | **Cell Signaling #7074S** |
| **Goat anti-Mouse IgG secondary antibody HRP** | **Mouse** | **1:1000** | **Stemgent #09-0039** |

**Supplementary Table S2: Used qPCR primers.**

| **Gene** | **Sequence** |
| --- | --- |
| **hOCT4** | **Fwd: GTGGAGGAAGCTGACAACAA**  **Rev: ATTCTCCAGGTTGCCTCTCA** |
| **hSOX2** | **Fwd: AGTCTCCAAGCGACGAAAAA**  **Rev: TTTCACGTTTGCAACTGTCC** |
| **hNANOG** | **Fwd: CCTGTGATTTGTGGGCCTG**  **Rev: GACAGTCTCCGTGTGAGGCAT** |
| **p21** | **Fwd: GGAAGACCATGTGGA**  **Rev: GGGGTTTGGAGTGGT** |
| **p53** | **Fwd: CAGGGCAGCTACGGTTTCC**  **Rev: CAGTTGGCAAAACATCTTGTTGAG** |
| **PCNA** | **Fwd: GACAAATGCTTGCTGACCTGG**  **Rev: TGAAGCCGAAACCAGCTAGA** |
| **hTERT** | **Fwd: CGGAAGAGTGTCTG**  **Rev: GGATGAAGCGGAGTCTGGA** |
| **KI67** | **Fwd: TCGTCCCAGTGGAAGAGTTG**  **Rev: CGACCCCGCTCCTTTTGAT** |
| **NPHS1** | **Fwd: GGAGGGTTGACTGGAAAAAGG**  **Rev: GCAACCACCACCACAATCG** |
| **NPHS2** | **Fwd: CTGTGAGTGGCTTCTTGTCCT**  **Rev: AGCAGATGTCCCAGTCGGAA** |
| **SYNPO** | **Fwd: CACCTGGAGAAGGTG**  **Rev: GACAGGTGCAGCCCA** |
| **CD2AP** | **Fwd: AACTCATGAAGCCCAGGACGA**  **Rev: CTGATCCAGATGCAGTTTCACTCAC** |
| **GAPDH** | **Fwd: GATTTGGTCGTATTGGGCGC**  **Rev: TTCCCGTTCTCAGCCTTGAC** |
| **MAPK8** | **Fwd: TGCAACCAACAGTAAGGACTTAC**  **Rev: TGAGTCAGCTGGGAAAAGGAC** |
| **STAT3** | **Fwd: TGATTTTAGCAGGATGGCCC**  **Rev: TTGGGAAGCTGTCACTGTAG** |
| **APOL1** | **Fwd: AATGAGGCCTGGAAC**  **Rev: TCAACCGAGGAAACT** |
| **Cycs** | **Fwd: CCGTCGGCGAGTACAACAAAG**  **Rev: CCAGCTCCACGTCCAAGAAGTAGT** |
| **Caspase 1** | **Fwd: CCCTGGTGTGGTGTGGTTTA**  **Rev:: CTCTTCACTTCCTGCCCACA** |
| **IL-6** | **Fwd: GGTACATCCTCGACGGCATCT**  **Rev: GTGCCTCTTTGCTGCTTTCAC** |
| **CLU** | **Fwd: ATTCATACGAGAAGGCGACG**  **Rev: CAGCGACCTGGAGGGATT** |
| **TGFβ** | **Fwd: TCAAGCAGAGTACACACAGC**  **Rev: TGCTGCTCCACTTTTAACTTG** |
| **NGAL** | **Fwd: GGTTTCATCCAGGATCGAGCAGG**  **Rev: ACAAAGATGGTCACGGTCTGCC** |
| **Gasdermin D** | **Fwd: CTGGAGTGCCTGGTGTTGTC**  **Rev: GCTGGTGGGCAAACACTCT** |

**Supplementary Figure S1: Uncropped Western blots**
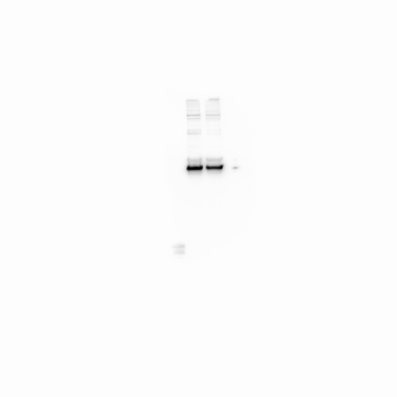
**.**

**E (Ladder)**

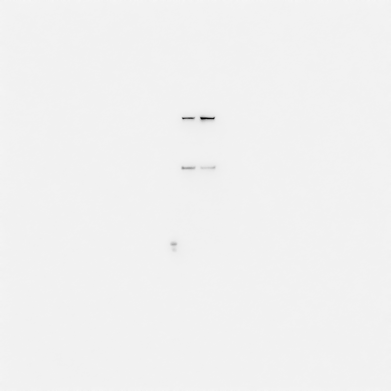

**A (α-ACTININ 4)**
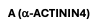
**4)**

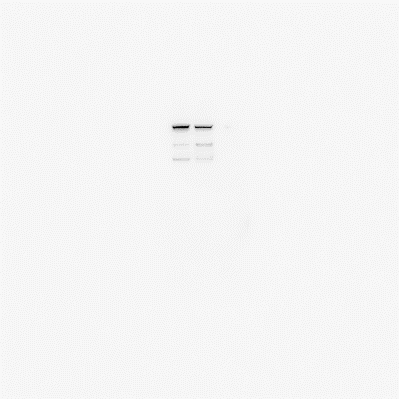

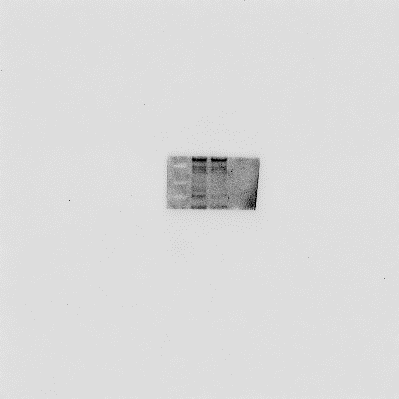

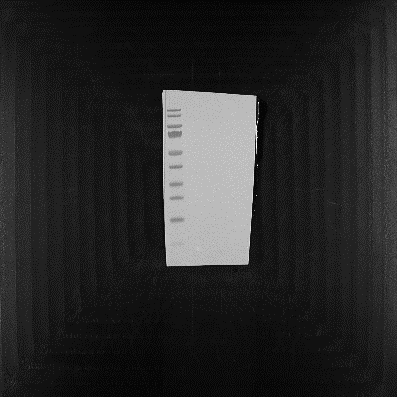

**D (GAPDH)**

**C (NPHS2)**

**B (NPHS1)**

**H (NPHS2)**

**G (NPHS1)**

**F (α-ACTININ 4)**
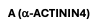
**4)**

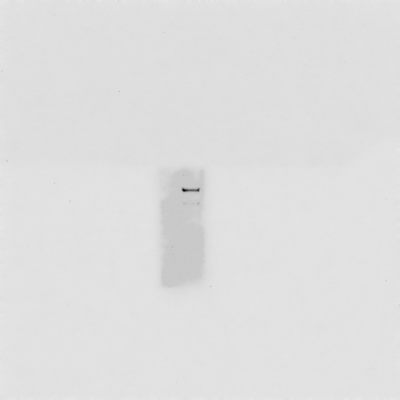

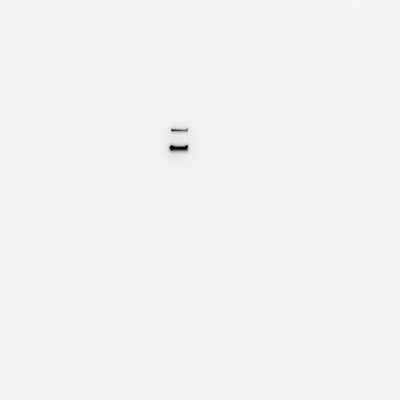

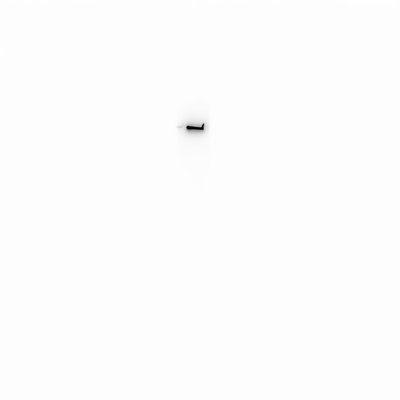

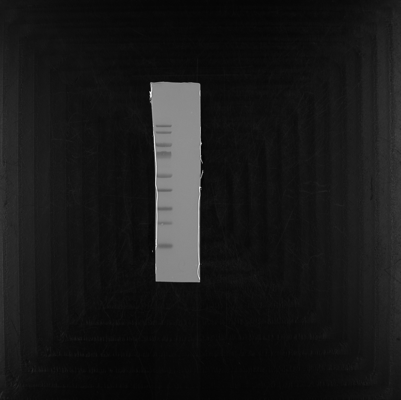

**I (GAPDH)**

**J (Ladder)**

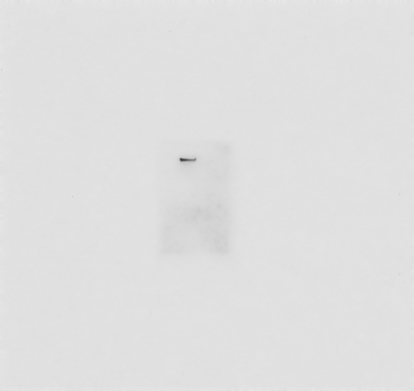

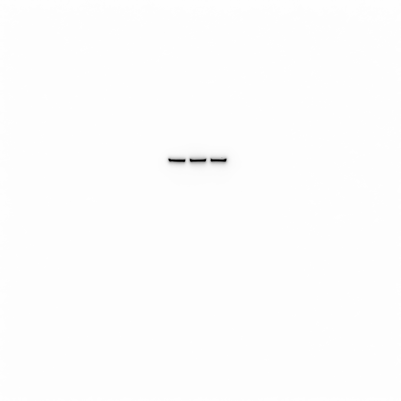

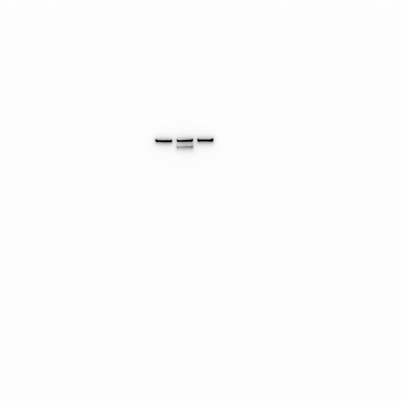

**K (α-ACTININ 4) ACTININ 4)**
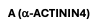

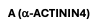
**)**

**L (p-STAT1)**
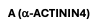
**)**

**O (GAPDH)**

**N (GAPDH)**

**M (APOL1)**

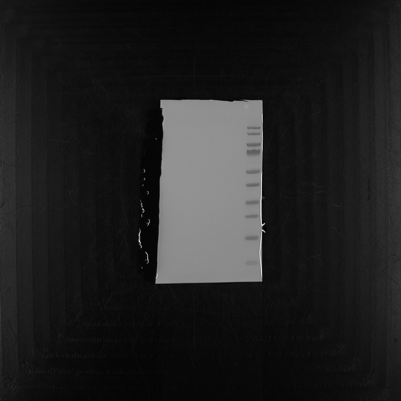

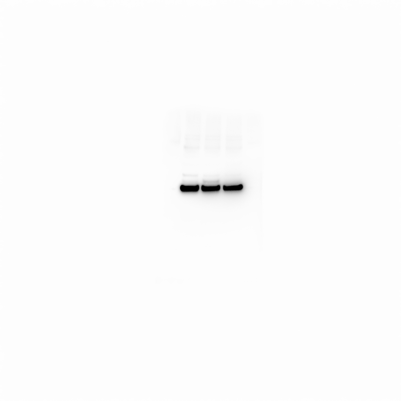

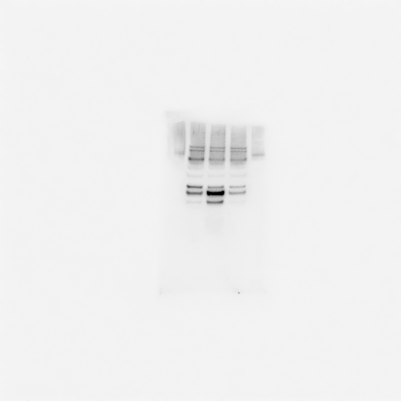

**Supplementary Figure S1: Uncropped western blot images.**

The detected proteins are presented from the left tot he right: A+F = $\alpha-ACTININ 4$, B+G = NPHS1, C+H = NPHS2, D+I = GAPDH. Supplementary figure E+J represents the ladder fort he western blot images. The loading scheme starting next to the ladder from the right for the western blots A-D is as following: UM30-OSN and UM30-OSN podocyte. The loading scheme for the western blots F-I also starting next to the ladder has only one sample the human immortal podocyte line (AB 8/13). All proteins show the predicted kDA size. Figure K-P represents the uncropped blots for Figure 5. The loading scheme starting next tot he ladder from the right like follows: UM30-OSN podocyte, UM30-OSN podocyte treated 24h with 100ng/ml IFNγ and UM30-OSN podocyte treated with 48h 1µM Baricitinib and 24h 100ng/ml IFNγ. Thereby the western blots K= $\alpha-ACTININ 4$, L=p-STAT1, M= APOL1, N= GAPDH and O= ladder. All proteins are detceted under their corresponding size.

**Supplementary Figure S2: Plasmidmaps.**

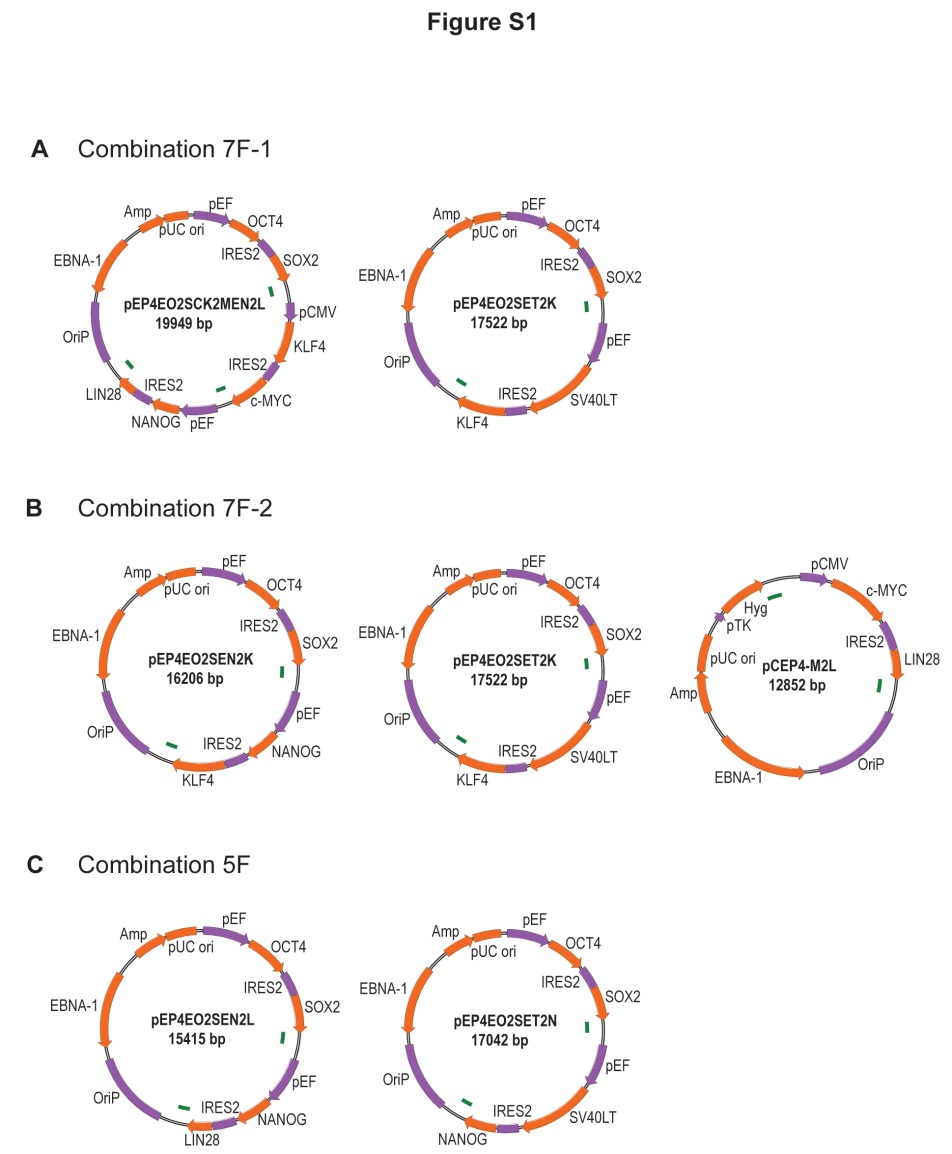

**Supplementary Figure S2: Plasmid maps.**

For the episomal reprogramming of the UM30-OSN the Combination 7F-1 plasmids were used.

**Supplementary Figure S3: Karyogram of UdRPC UM30-OSN and immortal cell line (AB 8/13).**

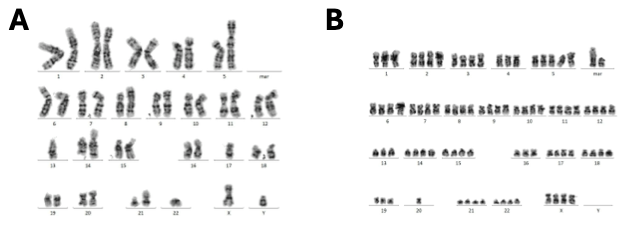

**Supplementary Figure S3: Karyogramm UM30-OSN and human immortal cell line (AB 8/13).**

(A) Karyogramm for the UM30-OSN cell line.The cytogentic.The Karyotyp of the UM30-OSN: 37-78,XYY,add(1)(p36)[3],add(3)(p25)[10],add[8](p21)[9],add(13)(p11)[5],add(14)(p11)[9],del(15)(q22)[3],add(16)(p13)[5],-17[11],der(21;22)(q10,q10)[22][cp24].
(B) Karyogramm for the human immortal podocyte line (AB 8/13). The Karyotype of the human immortal cell line(AB 8/13) is: 83-86,XXXX,-3,-13,-16,-19,-19,-20,+mar[21]/46,XX[1].

Both Karytypes were analyzed at the the Institute of Human Genetics of the University Hospital Düsseldorf.
